# An Atlas of Cortical Arealization Identifies Dynamic Molecular Signatures

**DOI:** 10.1101/2021.05.17.444528

**Authors:** Aparna Bhaduri, Carmen Sandoval-Espinosa, Marcos Otero-Garcia, Irene Oh, Raymund Yin, Ugomma C. Eze, Tomasz J. Nowakowski, Arnold R. Kriegstein

## Abstract

The human brain is subdivided into distinct anatomical structures. The neocortex, one of these structures, enables higher-order sensory, associative, and cognitive functions, and in turn encompasses dozens of distinct specialized cortical areas. Early morphogenetic gradients are known to establish an early blueprint for the specification of brain regions and cortical areas. Furthermore, recent studies have uncovered distinct transcriptomic signatures between opposing poles of the developing neocortex^1^. However, how early, broad developmental patterns result in finer and more discrete spatial differences across the adult human brain remains poorly understood^2^. Here, we use single-cell RNA-sequencing to profile ten major brain structures and six neocortical areas during peak neurogenesis and early gliogenesis. Our data reveal that distinct cell subtypes are predominantly brain-structure specific. Within the neocortex, we find that even early in the second trimester, a large number of genes are differentially expressed across distinct cortical areas in all cell types, including radial glia, the neural progenitors of the cortex. However, the abundance of areal transcriptomic signatures increases as radial glia differentiate into intermediate progenitor cells and ultimately give rise to excitatory neurons. Using an automated, multiplexed single-molecule fluorescent *in situ* hybridization (smFISH) approach, we validated the expression pattern of area-specific neuronal genes and also discover that laminar gene expression patterns are highly dynamic across cortical regions. Together, our data suggest that early cortical areal patterning is defined by strong, mutually exclusive frontal and occipital gene expression signatures, with resulting gradients giving rise to the specification of areas between these two poles throughout successive developmental timepoints.

## Introduction and Results

The specification of the brain into distinct regions has long been of interest to neuroscientists. Explaining how functionally distinct neural structures and cortical areas emerge bridges brain development with adult brain function. Understanding when brain regions acquire their unique features and how this specification occurs has broad implications for the study of human brain evolution, including species-specific developmental differences that may be responsible for the expansion of cortical areas such as the prefrontal cortex (PFC)^3^. It is also crucial for characterizing the pathology of neurodevelopmental and neuropsychiatric disorders, that often preferentially impact specific brain regions and/or cortical areas^4, 5^. Moreover, understanding brain patterning is essential for the accurate recapitulation of human brain development in *in vitro* models, including pluripotent stem cell-derived organoids.

Early patterning of the developing telencephalon is orchestrated by morphogenetic gradients of growth factors including bone morphogenetic proteins (Bmps), Wnts, Sonic hedgehog (Shh), and, most prominent in the cortex, fibroblast growth factor (FGF)^4, 6^. However, the molecular patterns that arise as a result later in development are less understood. Many of the seminal studies of cortical arealization took place prior to the widespread availability of next-generation sequencing and was performed in rodent models. Thus, numerous questions remain concerning human brain patterning and cortical arealization, such as the areal specificity of distinct cell populations, changes in gene expression throughout differentiation and maturation, and the processes by which these differences arise in the developing human brain.

To characterize the emergence of cellular diversity across major regions of the developing human brain and across cortical areas, we sequenced the transcriptome of single cells from distinct microdissected regions of developing human brain tissue samples along a ten-week window during the second gestational trimester, which encompasses peak stages of neurogenesis^7^. In this study, we refer to the subdivisions of the cerebrum and cerebellum as “regions”, and to subdivisions of the cerebral cortex as “areas”. We sampled cells from 10 distinct major forebrain, midbrain, and hindbrain regions from 13 individuals (Methods). Where available, we sampled: neocortex, proneocortex (cingulate), allocortex (hippocampus), claustrum, ganglionic eminences (GE), hypothalamus, midbrain, striatum, thalamus, and cerebellum (Fig 1A, STable 1). In addition, we sampled six neocortical areas from the same individuals: prefrontal (PFC), motor, somatosensory, parietal, temporal, and primary visual (V1) cortex. We used stringent quality control (QC) parameters (Methods), resulting in 698,820 high-quality cells for downstream analysis. We began by exploring the cell types and gene expression signatures in a whole brain dataset. Using an iterative clustering approach to mitigate batch effects (Methods), we characterized the broad cell types and states across the entire dataset. We found expected cell populations of the developing brain, including excitatory neurons, intermediate progenitor cells (IPCs), radial glia, mitotic cells, astrocytes, oligodendrocytes, inhibitory neurons, microglia, and vascular cells (including endothelial cells and pericytes) (Fig 1B). Unbiased clustering of cells, as well as their visual representation in UMAP space, were driven primarily by cell type rather than biological age, except for more mature glial populations, including astrocytes and oligodendrocytes, which were enriched in older samples (Fig 1B, SFig 1A).

**Figure 1.**
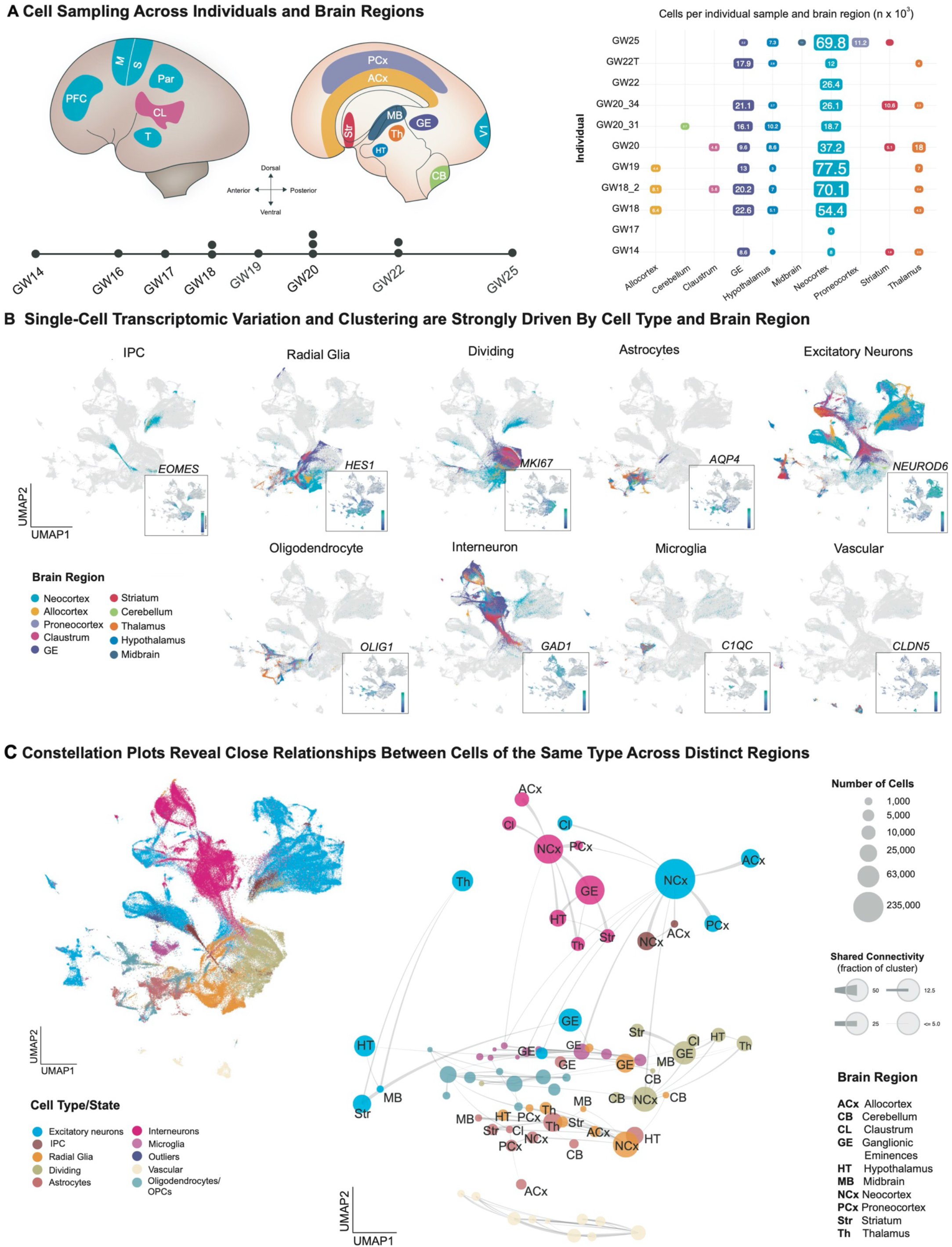
Single-cell analysis of gene expression signatures across regions of the developing human brain. **a)** Schematic on the left shows the anatomical brain regions sampled for this study. The timeline below highlights the number of individuals sampled at each gestational week. The matrix on the right shows the final count (after QC) of cells from each individual distributed across regions sampled. **b)** Single cells from all brain regions sampled are represented in UMAP space. Cells are color-coded by their region of origin. Insets show the expression profile of canonical genes representative of each identity. **c)** Top left panel: Distribution of cell types/states in UMAP space. Constellation plot of cells grouped by type/state and brain region highlights the interplay between cell type (node color) and regional identity (node label). Nodes are scaled proportionally to the number of cells in each group. Edge thickness at each end represents the fraction of cells within a group with neighbors in the opposite group. Node color corresponds to cell type/state; node label corresponds to the brain region from which cells were sampled.

We next sought to identify marker genes specific to or significantly enriched in distinct cell populations in each brain region. We found genes that were region-specific across all cell types, as well as genes that were region-specific for individual cell types (STables 2 and 3). We detected previously described markers of brain regions, including FOXG1 (cortex)^8^, ZIC2 (cerebellum, also observed in the neocortex)^9, 10^, and NRP1 (allocortex)^11^,(SFig 1B), as well as other region-specific genes, many of which are transcription factors (STable 2). We also identified numerous cell type and structure-specific transcription factors, including OTX2, GATA3, LHX9 and PAX3. The distribution of cell types across regions was as expected, with progenitor and differentiated cell types in each region (SFig 1C), leading us to ask whether brain region or cell type is a stronger component of regional identity during the second trimester. At earlier developmental timepoints, we and others have noted that regional signatures are not broadly pervasive and do not yet reflect area-specific identities of unique brain substructures^12,13^. To perform this analysis, we first used hierarchical clustering to group individual cell type subclusters. As expected, cluster branches were primarily organized by cell type, validating our annotation approach and highlighting the robustness of cell type in driving cluster similarity. However, by quantifying the proportion of cells from each region contributing to each cluster, it became apparent that the majority (115 of 192) of clusters were strongly enriched for a single brain region (i.e. cortex, thalamus, etc.) or for several related regions (e.g. forebrain) (SFig 1D). A small number of inhibitory and excitatory neuron clusters bridged across regions, while a larger proportion of glial and vascular cells could be identified across brain regions, indicating a strong regional signature for each individual cell type.

To further interrogate the interplay between cell type and brain region identity, we generated constellation plots using cells’ combined brain region and cell type annotations. Constellation plots are a powerful tool to visualize the relationships between groups of cells, by quantifying the proportion of cells in a group or cluster with an above-threshold number of nearest neighbors in other groups, while preserving the UMAP topography of the dataset (Methods). We find that across the whole brain, cell type is the primary source of segregation (Fig 1C). However, in certain cases, such as the GE, cells of distinct types from a common region are drawn together in UMAP space, suggesting that the regional identity conferred upon groups of cells of different types can also be a strong source of variation (Fig 1C). A heatmap of area-specific gene score enrichments (Methods) shows that some region-specific genes are present across multiple cell types within a given region. This suggests that some regional gene expression signatures are highly penetrant across cell types. Regionalization is stronger in glial populations at the developmental timepoints we studied (SFig 1E). Thus, we identify strong, cell-type independent regional signatures, as well as signatures that are restricted to specific cell types (STable 3).

The neocortex, allocortex, and proneocortex are evolutionarily closely related, and physically proximal^14^. However, we sought to identify distinct regional gene expression programs among these three closely related regions by co-clustering these samples in isolation from the rest of the brain (SFig 2A). We observed extensive integration of cells from the neocortex and allocortex, but segregation of the proneocortex which was sampled at a later timepoint (SFig 2A). We set out to identify the similarities and differences between cell-type and regional expression signatures between these three cortical structures. Using differential gene expression in each excitatory lineage cell type (STable 4), we again used a gene score annotation paired with hierarchical clustering (SFig 2B). Surprisingly, even within these closely related cortical structures, region was still the primary driving force, and again, regional signatures bridged multiple cell types (SFig 2C-E). These analyses indicate that during the second trimester of development, regional signatures are sufficiently established to distinguish cells across brain structures, with some of these signatures extending beyond an individual cell type. We provide analysis of these region-specific gene signatures, including a cell-type specific analysis (STable 3, SFig 2C-E).

The neocortex is made up of dozens of functional areas that specialize in an astounding range of cognitive processes, from sensory perception to reasoning, decision-making and language^15^. Longstanding, juxtaposed hypotheses propose the existence of either a cortical *protomap*^*16*^, where the areal identity of cortical progenitors, and consequently that of their progeny, is cell-intrinsic and genetically predetermined, or a *protocortex*, where newborn neurons are not areally specified until extrinsic signals, such as those from thalamocortical afferents, reach the developing cortex^17^. Single-cell RNA-sequencing has the power to deconvolute areal gene expression signatures and therefore may provide a means to test these models. Recent work has shown that while neurons are distinct between V1 and PFC soon after their birth^1^, other cell types do not show such large area-specific differences. Studies in the adult mouse have additionally shown that neuronal cell types of the anterior lateral motor cortex (ALM) and V1, are transcriptionally distinct from each other^2^, but that denser sampling of areas between the ALM and V1 reveals a gradient-like transition between cell type profiles^18^. We sought to expand upon these findings by profiling single cells from distinct cortical areas in order to clarify how and when distinctions between areas begin to emerge. To do so, we performed single-cell RNA-sequencing of six cortical regions subdissected from the samples described above, yielding 387,141 high-quality cells, after filtering (Methods) (SFig 3A-B), for subsequent analysis.

We applied the same iterative clustering and annotation methods used for our analysis of the whole brain. We found expected cell types, including Cajal-Retzius neurons, dividing cells, excitatory neurons, inhibitory neurons, IPCs, microglia, oligodendrocyte precursor cells, radial glia/astrocytes, and vascular cells (Fig 2A, STables 5 and 6). Hierarchical clustering of 138 neocortical clusters grouped cells by cell type (Fig 2B). showed that most clusters (104/138) are composed of cells from multiple cortical areas.

**Figure 2.**
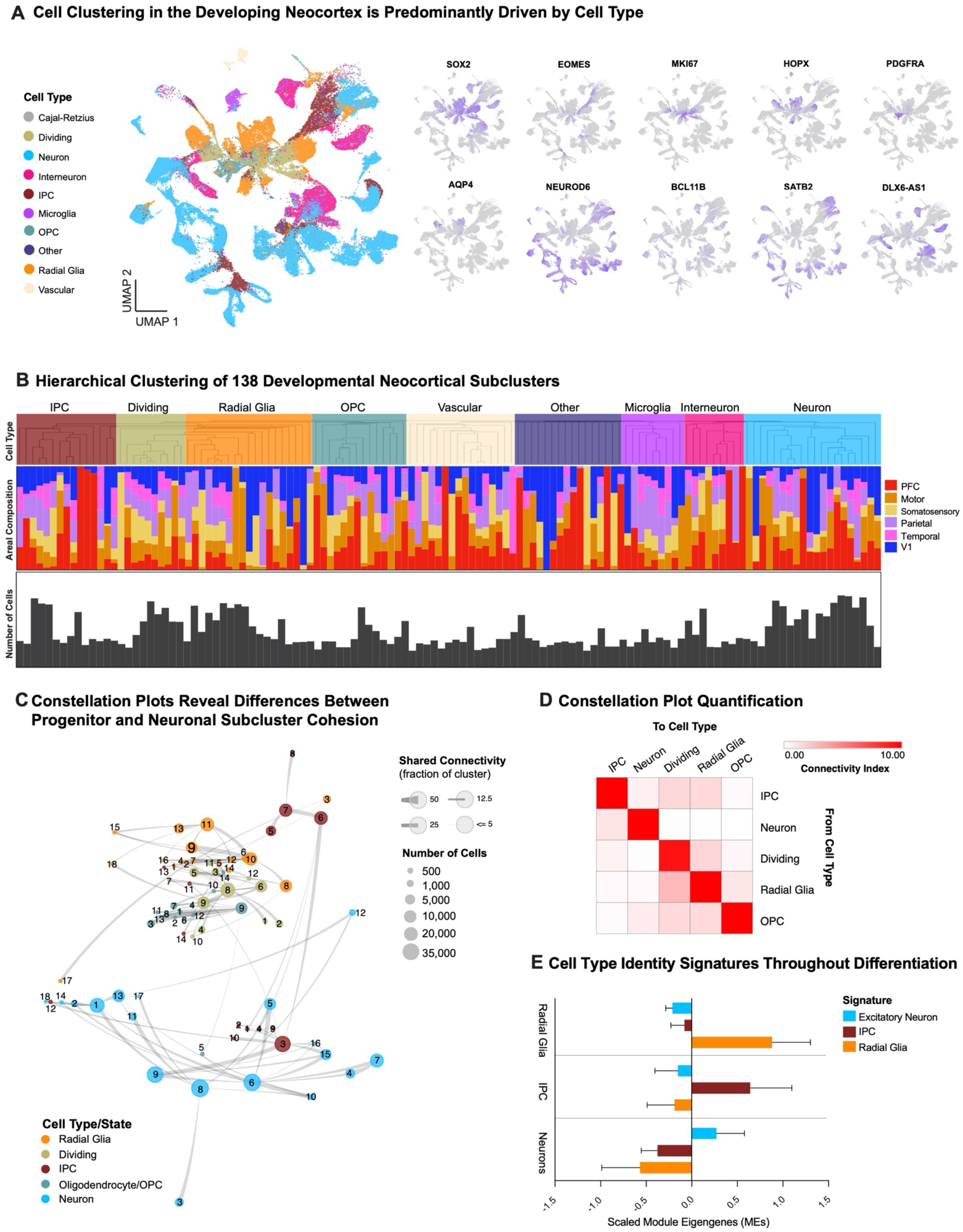
Cell Types in the Developing Human Neocortex Across Cortical Areas. **a)** Single-cells from the neocortex represented in UMAP space. Cells in the left-most UMAP plot are color-coded by cell type/state annotation. Feature plots on the right depict the expression pattern of major cell population markers (SOX2 – radial glia, EOMES – IPC, MKI67 – dividing cells, HOPX – outer radial glia, PDGFRA – OPC/Oligodendrocyte, AQP4 – astrocytes, BCL11B – deep layer excitatory neurons, SATB2 – supererficial layer excitatory neurons, NEUROD6 – broad excitatory neurons, DLX6-AS1 – inhibitory neurons). **b)** Hierarchical clustering of 138 neocortical clusters based on the Pearson correlations of cluster marker expression profiles across all neocortical stages sampled. Branches are color-coded by the major cell type assigned to each cluster. Histograms below show the fraction of cells from each area contributing to a cluster. Bottom bar chart shows the relative number of cells in each cluster (log2-transformed numbers,0 to 20). **c)** Constellation plot shows the relationships between clusters of the excitatory lineage and oligodendrocytes, highlighting the strong relationships between clusters of the same cell type and lineage, but not between those of different groups. Nodes are scaled proportionally to the number of cells in each group. Edge thickness at each end represents the fraction of cells within a group with neighbors in the opposing group. Nodes are colored by cell type/state, and labelled by the brain region from which cells were sampled **d)** Constellation plot quantification, with ‘towards cell type’ connectivity as columns and ‘from cell type’ connectivity as rows. The connectivity index integrates the number of connections between two distinct cell types, as well as the average fraction of contributing cells from each cluster. **e)** Differential expression was performed at the cell type level between radial glia, IPCs, and excitatory neurons. Each set of cell-type-enriched genes was used to create a network and scaled module eigengenes were calculated for each cell type.

To explore how distinct each cluster was from others of the same cell type or lineage, we again used a constellation plot approach (Fig 2C). We found strong connectivity between clusters of the same cell type, suggesting that borders between clusters are fluid. We also found some connectivity, albeit to a lesser degree, between cells from the excitatory lineage. To quantify intra-cell type and inter-cell type connectivity, we calculated the magnitude of transcriptional proximity between nodes (Methods), and found, not surprisingly, that clusters of each cell type connected most strongly to each other (Fig 2D). However, we also found that when comparing inter-cell type connectivity, IPC subclusters connected much more strongly with excitatory neurons than with radial glia subclusters (Fig 2D).

We then defined gene signatures characteristic of radial glia, IPCs, and excitatory neurons in the neocortex using a differential gene expression approach (Methods). These signatures (STable 7) were scored using a module-eigengene calculation (Methods) across the major excitatory lineage cell types (radial glia, IPC, excitatory neurons). We found that radial glia had the highest up-regulation of the progenitor signature, but lower down-regulation of the IPC and neuronal signatures. In contrast, excitatory neurons had the lowest up-regulation of the neuronal signature but strongest downregulation of other programs, perhaps reflecting lack of neuronal maturation. Together with the differences in connectivity strength between cell types seen in the constellation plots, this suggests that there is a strong break between radial glia and excitatory neuron identities, but interestingly, that IPCs track more closely with neurons than radial glia, despite their progenitor nature. These differences in cell type gene signature strength and cell type connectivity reveal a cascading differentiation program along the excitatory lineage in the neocortex.

In addition to systematically exploring gene expression signatures between excitatory lineage cells along the axis of differentiation, we sought to understand how cortical area influences the identity of cells in the developing cortex, as well as the relationships between cell clusters. We first validated our cortical sub-dissections by visualizing the gene expression of NR2F1, a posterior-high to anterior-low expression gradient marker, ^19^ in our dataset (Fig 3A) and previously described cortical area specific genes (SFig 3C). Next, we built constellation plots using a different cell grouping approach, with each group corresponding to a specific area and cell type (Fig 3B). Several striking patterns emerged. First, we noted that radial glia nodes were connected primarily to other radial glia, while IPCs and excitatory neurons were frequently mutually connected to one another (Fig 3B - C). This break between radial glia versus IPCs and glutamatergic neurons suggests large differences in cell type signature and possibly also areal signatures between the groups. Radial glia from distinct cortical areas were highly interrelated and formed a tight, insular network, with sparse edges to their descendant cell types, IPCs and pyramidal cells (Fig 3B). In contrast, excitatory neurons from distinct cortical areas were less interrelated and formed a looser network when compared to radial glia. Of note, neuronal nodes of different areas show robust area-specific transcriptional proximity to their IPC counterparts, suggesting that some degree of the areal specification seen in neurons is already present at the IPC stage (Fig 3B). We did not find edges between PFC and V1 cell type nodes (Fig 3C), suggesting a model of strong mutual exclusivity between these two gene expression programs. These patterns are persistent when including cell subtype annotations, as well as when analyzing individual developmental stages separately (SFig 4A – H). This was consistent with previous observations of early specification of PFC and V1 identity, so we additionally quantified the number of differentially expressed genes across each cell type by area (Fig 3D, SFig 3D). Consistent with prior observations^1^, the specificity of neuronal areal markers was significantly higher compared to radial glia (Fig 3D) but surprisingly, more genes were differentially expressed in radial glia than in neurons (Fig 3D, STable 8). These two observations indicate that markers of areal identity are already detectable in radial glia but become more pronounced as differentiation proceeds, and importantly, there is a significant overlap between the area-specific genes we describe here and those in previous studies (SFig 3E).To further interrogate the relationship between differentiation and the transformation of regional signatures across development, we inferred lineage trajectories across cells using RNA-velocity^20, 21^. This uncovered strong trajectories along known differentiation gradients, primarily between radial glia, IPCs, and excitatory neurons (SFig 5A). For each individual cortical area, we identified the most dynamic genes across the differentiation cascade as the top loading velocity genes (SFig 5B). Hierarchical clustering of the underlying counts values of these genes across excitatory lineage cells depicted an expected segregation by cell type, and within the excitatory neuron populations, showed small clusters of unique enriched genes across all areas. The gene enrichment in excitatory neuron groups, but not in radial glia or IPC populations led us to ask how areal signatures might change during differentiation. We defined areal signatures of excitatory neurons as gene networks and evaluated their representation in radial glia across cortical areas by calculating their module eigengene scores (SFig 5C). We found an early strong binary V1 expression, while PFC gene signature emerged only at later developmental timepoints (SFig 5C).

**Figure 3.**
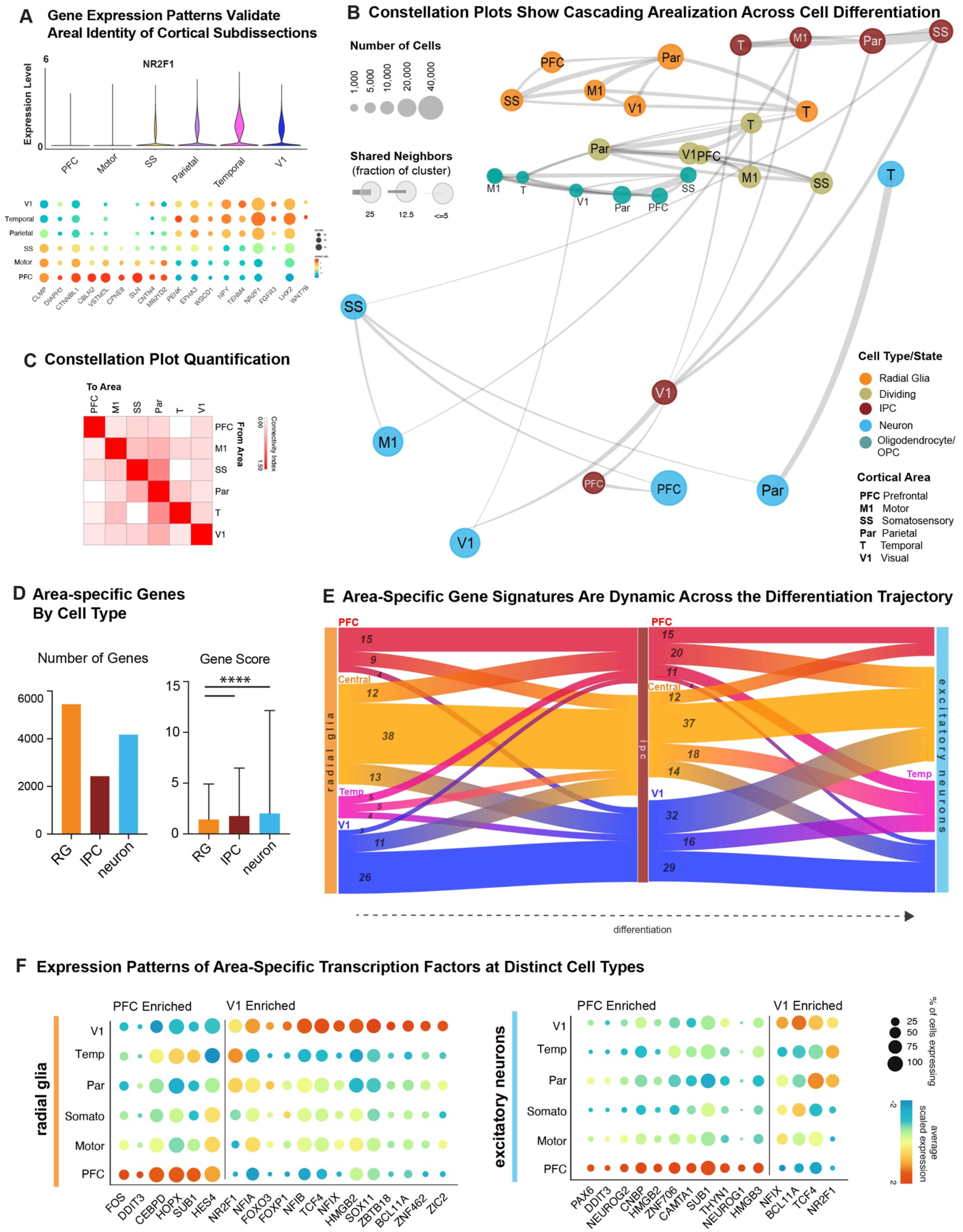
Cortical Area-Specific Gene Signatures. **a)** Top: Violin plots show the expression of previously described posterior-high to anterior-low gradient marker gene NR2F1 across all neocortical cells grouped by area. Bottom: Dot plot shows the expression of a representative panel of previously reported areally enriched genes across all neocortical cells grouped by area. Expression profiles validate the areal identity of the cortical subdissections used in this study, **b)** Constellation plots of excitatory lineage and oligodendrocyte lineage grouped by cortical area highlight similarities between groups of frontal and occipital cortical areas. Each dot is scaled proportionally to the number of cells represented by that analysis. The thickness of the connecting line on each end represents the fraction of cells within each group with neighbors in connected groups. Dot color represents cell type while text over the dot marks cortical area. **c)** Quantification of the constellation plots, with ‘towards area’ in columns and ‘from area’ in rows. The connectivity index from white to red integrates the number of connections between two cell types as well as the average fraction of cells from each cluster contributing to each connection. **d)** Quantification of the number of differentially expressed areal genes from each cell type, using a union of all genes as calculated across each individual in the dataset. The gene score, an integration of fold change and specificity is quantified in the lower graph, with mean plus standard deviation shown. Neurons vs radial glia p-value = 0.000011; IPC vs radial glia p-value = 0.000435 (one-sided t-test). For both analyses, n= 5446 (radial glia), 2426 (IPCs), 4170 (Neurons). **e)** Sankey plots show the proportion of area-specific gene groups shared between cell types and cortical areas. The number on each block line indicates how many genes are represented by that stream as the stream sizes are not to scale. The “central” area encompasses motor, parietal, and somatosensory areas. **f)** Dot plots quantify a subset of transcription factors enriched in PFC or V1 radial glia (left) and excitatory neurons (right) relative to other cortical areas. Enrichment can occur via an increase in the number of cells expressing a given gene, an increase in the average expression level of expressing cells, or both.

We sought to explore how areal gene signatures change throughout neuronal differentiation by constructing Sankey diagrams to display the overlap between area-specific markers of each cell type. We found that area-specific gene expression signatures change substantially across cell types, with small numbers of areal markers preserved throughout the differentiation trajectory (Fig 3E). This is in stark contrast to the pervasive quality of regional signatures which were present across distinct cell types across the whole brain. This suggests that cell type identity is a much stronger contributor to a cell’s transcriptional profile than cortical areal identity at these developmental stages.

Within each set of areal marker genes, we identified several transcription factors that were robustly enriched in cells of a specific area relative to all other areas, as well as TFs with a broader frontal or caudal enrichment. These transcription factors are shown in a dot plot format^22-25^ to clearly depict both expression level and fraction of cells across cortical areas (Fig 3F). These observations suggest that each cortical area is anchored with core transcriptional programs, some of which might persist across cell type boundaries (Fig 3F, SFig 6A). A subset of marker TFs show consistent area specificity throughout the entire developmental window we studied, i.e. at early, middle and late second trimester (SFig 6A). We detected TFs with established roles in arealization, such as NR2F1, which confers positional identity across the rostro-caudal axis^19^, and BCL11A, which interacts with NR2F1 and was recently shown to repress motor identity in the cortex^26^. Both genes are also implicated in neurodevelopmental disease^27, 28^. We also detected TFs that, to our knowledge, have not been previously described in the context of cortical arealization. In V1, these markers include members of the Nuclear Factor I family, NFIA, NFIB and NFIX. These genes are important regulators of brain development and have been implicated in developmental disorders including macrocephaly and severe cognitive impairment^29^. They also include ZBTB18/RP58, a transcription factor involved in neuron differentiation and cortical migration and a putative driver of brain expansion^30, 31^. In the PFC, area-specific TFs we identified include members of the HMGB family, HMGB2 and HMGB3, which are differentially expressed by neural stem cells at distinct stages of development^32^ and are thought to be key regulators of differentiation. Of note, HMGB3 mutations can result in severe microcephaly. We also found the TFs NEUROG1 and NEUROG2 to be upregulated in PFC neurons. As in V1, while these genes have been found to be important regulators of neuronal differentiation, they have not been previously implicated or studied in the process of cortical arealization.

In addition to exploring differences in areal gene signatures at the cell type level, we investigated these signatures change throughout development. Consistent with proposed models of extensive transcriptional remodeling during the second and third trimesters^33^, we observed that while area-specific gene signatures are composed of significant and specific marker genes, they also changed substantially throughout the early, middle and late second trimester (SFig 6B - C). Concordantly, we only found a small overlap of area-specific gene signatures, and low cluster correspondence, between this dataset and that of the adult brain (SFig 7A-C). Our analysis indicates that after the second trimester, neurons in the neocortex remain largely immature, and that while the extreme rostral and caudal identities are already determined in early progenitor populations, further areal specification is refined at later time-points and may depend on sensory inputs to determine terminal identity. We thus find strong evidence for a partial early cortical protomap, which is then further refined throughout development as proposed by the protocortex model.

Our single-cell data uncovers a large diversity of cell types and transcriptional profiles across six areas of the developing human cortex. We selected candidate markers of excitatory neuron clusters that were enriched in one or more sampled areas for validation by multiplexed single-molecule fluorescent *in situ* hybridization (smFISH) (Fig 4A). We used the Rebus Esper spatial omics platform (Rebus Biosystems, Inc), a novel, automated system based on synthetic aperture optics (SAO) imaging. We quantified the expression level of 31 RNA transcripts per tissue section at four cortical regions from a GW20 sample (Fig 4B). This resulted in data from an additional 608,960 cells, enabling us to visualize marker gene expression levels, their spatial distributions, and co-expression patterns. We used DAPI staining along with kernel density expression (KDE) plots^34^ of canonical cell type markers genes *SOX2, SATB2*, and *BCL11B* to identify the ventricular zone and cortical plate (Fig 4C, SFig 8 - 9). Additionally, we confirmed the previously described areal pattern dynamics between the neuronal genes SATB2 and BCL11B, which are co-expressed in frontal regions but mutually exclusive in occipital regions^1^ (Fig 4C). These spatial datasets are available for exploration and analysis at kriegsteinlab.ucsf.edu/datasets/arealization (STables 9 - 12).

**Figure 4.**
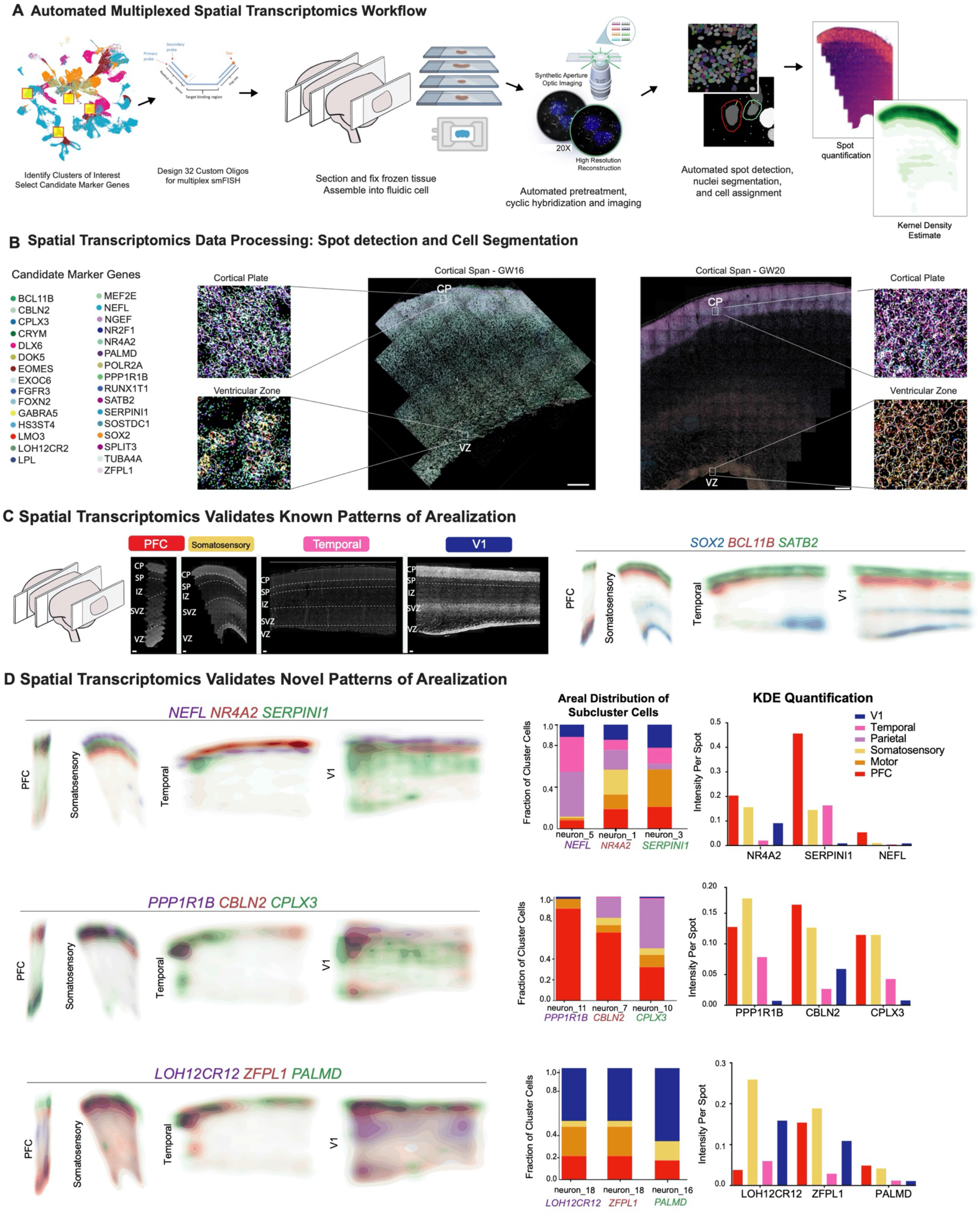
Spatial RNA Analysis Identifies Distinct Spatial Patterns of Area Specific Clusters. **a)** Automated Rebus Esper spatial RNA Transcriptomics workflow used to validate the expression patterns of candidate marker genes *in situ* across four distinct cortical areas. 31 relevant candidate marker genes of neuronal subclusters were first selected from our analysis. 22-96 custom oligos were designed for each RNA target. Tissue blocks from 4 cortical areas of a GW20 and a GW16 sample were sectioned (7-10um) to coverslips and mounted into a fluidic chamber, where iterative single-molecule fluorescent *in situ* hybridization (smFISH) was performed in batches of 3 genes at a time. RNA molecules were quantified and assigned to individual cells via automated spot detection and nuclei segmentation. These spots were then analyzed either by cell assignment or using their overall distribution pattern. **b)** Representative merged images of smFISH for 31 candidate marker genes in a GW16 (left) and GW20 (right) somatosensory cortex section. Zoomed in images of the ventricular zone (left) and cortical plate (right). White circles indicate segmented nuclei. Scale bar = 444 μM **c)** Top left: Nucleus staining outlines tissue architecture, with the ventricular zone at the bottom and the cortical plate at the top. Top right: kernel density estimate (KDE) plots of positive control genes. SOX2 marks radial glia and the ventricular zone, while SATB2 and BCL11B mark the cortical plate. As previously described, SATB2 and BCL11B are co-expressed in frontal regions, but are mutually exclusive in occipital regions. Scale bar = 444 μM **d)** KDE plots of neuronal genes of interest. Genes were chosen as candidate markers for specific neuronal subclusters. Clusters being explored are named below the histogram each gene marker for this cluster is below its name. Stacked histograms show the expected ratio of clusters as a fraction of total composition. To the far right in each row, the quantification of the KDE plots is shown as intensity divided by the number of spots in order to reflect both the intensity of signal but also the pervasiveness of the marker to not artificially bias the analysis by examples of rare but intense signal. We see strong correspondence between the predicted spatial distribution of clusters and the signal in our spatial RNA analysis.

Across all areas, we explored novel candidate markers of subpopulations, including genes that are also subplate markers (*NEFL, SERPINI1*). For these KDE plots, we also plotted the canonical subplate marker *NR4A2*. All three markers could be found at roughly equal intensities across cells in the PFC, somatosensory, temporal and V1 cortex, but their spatial distribution relative to one another changed substantially (Fig 4D). These genes were co-expressed in the PFC, but were mutually exclusive across all other regions. However, in the somatosensory cortex, we found these markers to be expressed in upper cortical layers rather than in the subplate. We similarly identified three frontally-enriched marker genes, *PPP1R1B, CBLN2*, and *CPLX3*, whose quantification also revealed higher signal in the PFC and somatosensory tissue sections after normalization (Fig 4D). Caudally, we observed higher intensities of *LOH12CR12, ZFPL1*, and *PALMD* (Fig 4D). In both sets of markers, we found striking differences in the laminar distribution of gene expression, suggesting that not only are gene expression levels genes different across the cortical rostro-caudal axis, but that laminar cell type distribution also changes (SFig 10A). While this observation may be reflective of differences in maturation states across the developing cortex, cell types may express genes in a different manner across distinct cortical areas.

We leveraged cell segmentation aspects of our spatial transcriptomics analysis to investigate how the co-expression patterns of the genes queried in this experiment changed across areas. Transcripts detected with smFISH were automatically assigned to nuclei by proximity with the Rebus Esper imaging processing software. We calculated co-expression relationships between single cells to generate networks show the frequency of two genes expressed the same cell (SFig 10B). The resulting networks highlight that the most stringent markers of areal identity are binary, i.e., they are either included or excluded from the gene network. In most cases, however, we found remodeled co-expression patterns across cortical areas rather than elimination or inclusion of a gene from the network. Even when with using all 31 genes to construct the networks, we see substantial co-expression remodeling across cortical areas. We repeated this spatial analysis in a second individual (GW16), and validated some of the area specific aspects of gene expression while also continuing to observe dynamic laminar redistribution across cortical areas (SFig 11 – 14, STable 13 – 16). In sum, these data provide *in situ* evidence that reinforces our observation from single-cell RNA seq analysis that there exist area-specific and regional-specific cell populations.

In this study, we performed an in-depth characterization of the transcriptomic patterns and gene expression dynamics of the developing human brain during mid-gestation. We first analyzed molecular signatures of brain regionalization across ten major brain regions, and subsequently focused on the neocortex and its specification into distinct areas. Our results provide a granular understanding of the gene expression signatures of distinct cell types across neocortical areas, and at eight different timepoints throughout the second trimester of development. We find that across major brain structures, regional identity is highly pervasive among distinct cell types. In contrast, areal identity in the neocortex is highly specific and restricted to individual cell types. Furthermore, we find that that in addition to cell type identity, the developmental stage of cells (i.e. gestational week) is a strong determinant of gene expression signature composition. Together, these observations suggest that the dynamics of area-specific gene expression signatures are surprisingly fast moving and cell type-specific (Fig 5 C, D). This is in contrast with previous models of areal patterning, where gene expression programs have been generally assumed to be persistent once established at a given region.

**Figure 5.**
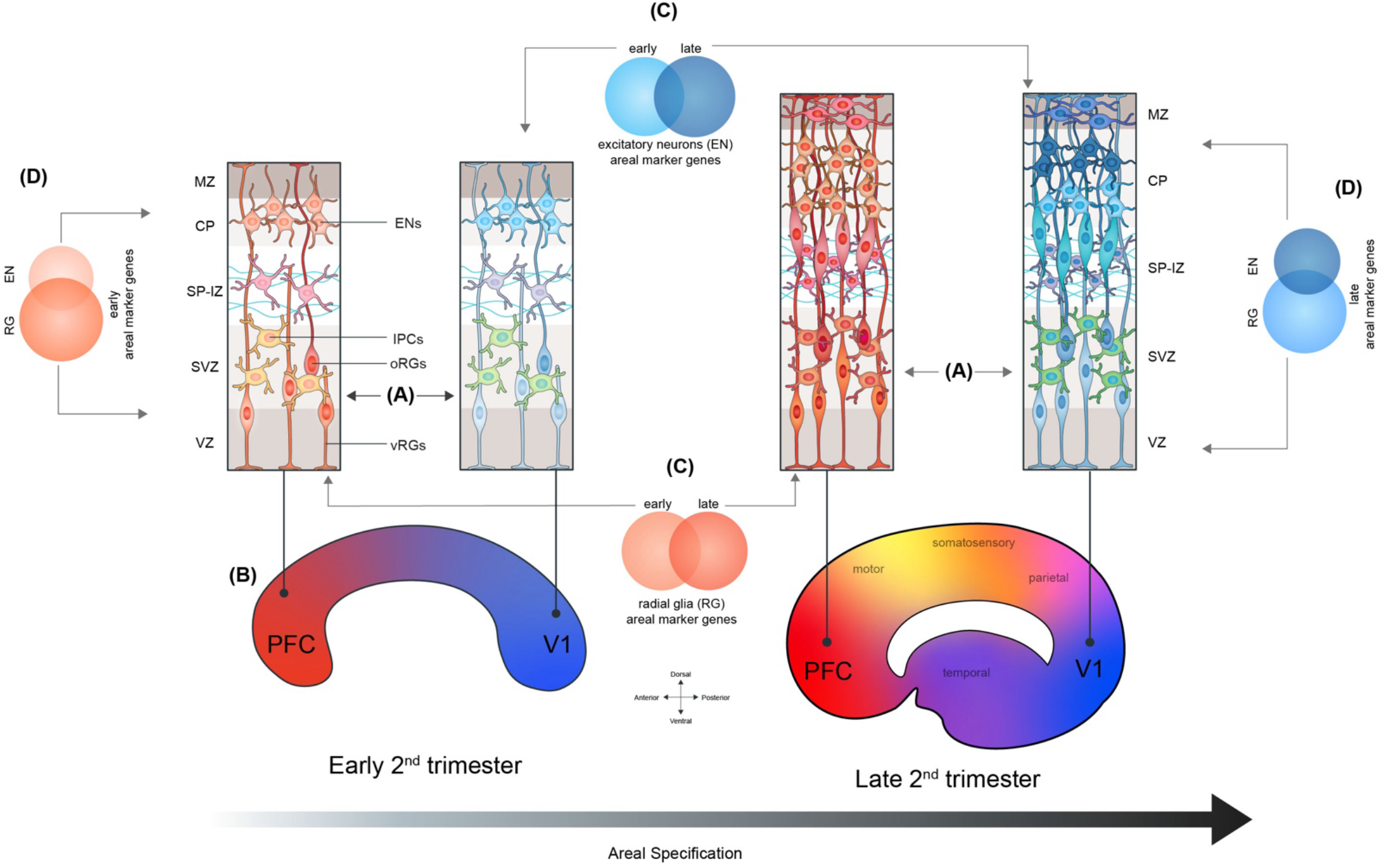
Summary Schematic of Proposed Model of Arealization. In this study, we find that in addition to their neuronal progeny, radial glia from distinct cortical areas are already distinct from each other (A). At early second trimester timepoints, the strongest contrast is seen radial glia at the frontal and occipital poles of the neocortex (B). The gene expression signatures of cells at different cortical areas are highly dynamic across developmental time (C) and across the differentiation axis (RG → IPC → excitatory neuron) (D). As differentiation progresses, these dynamic gene signatures give rise to other major cortical areas, and further refinement occurs, likely as sensory input to the cortex begins to take place. (E)

We find strong evidence for the presence of a partial early cortical protomap between cell populations, including progenitors, at the frontal and occipital poles of the neocortex (Fig 5 A, B). We see evidence of transcriptional regulation programs that may prime more differentiated and mature cells to acquire either a rostral or caudal identity. For example, even though progenitor clusters in the neocortex show little molecular diversity in relation to the multiple cortical regions that will eventually emerge, we observe strong specification of PFC and V1 molecular identity among cells of this cell type. In a previous study, we noted that radial glia were characterized by a small number of transcriptional differences that cascade into strong area-specific excitatory neurons^1^. Here, we present an analysis of nearly 400,000 cells compared to the 4,000 cells in the previous study. The analysis of a much greater number of cells and more cortical areas enables us to uncover the strong difference between PFC and V1 radial glia, while confirming that glutamatergic neurons are even more distinct between cortical areas. Our data suggests that cells located in between the prefrontal and occipital poles are less specified towards a particular areal identity, an observation that is more consistent with the *protocortex* hypothesis. Together, these observations suggest that the expression gradients that establish early neocortical areal patterning may be propagated throughout development through transcriptional or epigenetic memory.

Importantly, we find strikingly distinct spatial distributions of excitatory neuronal marker genes across areas *in situ*, as well as gene co-expression patterns unique to specific cortical areas. These differences may help explain the distinct morphological features of cortical areas, as well as the distinct connectivity patterns seen between areas and from different areas to other parts of the brain.

Characterizing the dynamic diversity of cell populations throughout the development of a structure as complex as the brain involves disentangling multiple axes of variation. The data we present here provides a spatially and temporally detailed molecular atlas of human brain and neocortex specification upon which future experimental characterizations can expand. These findings can additionally improve our understanding and development of *in vitro* models of cortical formation, including those used in the study of neurodevelopmental disorders.

## Acknowledgements

The authors thank Shaohui Wang, William Walantus, Madeline G. Andrews, Lakshmi Subramanian, and members of the A.R.K. laboratory for providing resources, technical help, and valuable discussions. We also thank Brian M. Filarsky for providing extensive computational guidance and support, and Pablo E. García and the Single-Cell Genomics Program at the Chan Zuckerberg Initiative (CZI) for their assistance in building and hosting the single-cell browsers that accompany this manuscript. Single-cell RNA sequencing data has been deposited at the NeMO archive under dbGAP restricted access. Some primary human tissue was obtained from the Human Developmental Biology Resource (HDBR), with special thanks to S. Lisgo and M. Crosier. This study was supported by NIH award U01MH114825 to A.R.K., F31NS118934 to C.S.E., and F32NS103266, K99NS111731, and L’Oreal For Women in Science Award through the AAAS to A.B. The data analyzed in this study were produced through the Brain Initiative Cell Census Network (BICCN: RRID:SCR_015820) and deposited in the NeMO Archive (RRID:SCR_002001). All code and datasets used in this study, along with single-cell and spatial transcriptomics browsers are available at kriegsteinlab.ucsf.edu/datasets/arealization.

## Author Contributions

A.B., C.S.E., T.J.N. and A.R.K. designed the study and analysis. Experiments were performed by A.B., C.S.E., M.O and T.J.N. Data analysis was performed by A.B., C.S.E. and U.C.E. The study was supervised by A.B. and A.R.K. This manuscript was prepared by A.B. and C.S.E. with input from all authors. The corresponding authors certify that A.B and C.S.E should be considered first-authors to all academic and professional effects, and can be indicated as such on their respective publication lists.

## Online Methods

### Sample Acquisition

De-identified tissue samples were collected with previous patient consent in strict observance of the legal and institutional ethical regulations. Protocols were approved by the Human Gamete, Embryo, and Stem Cell Research Committee (institutional review board) at the University of California, San Francisco. Two sets of samples included twins: GW20_31 and GW20_34; GW22 and GW22T.

### Single-cell RNA Sequencing Capture and Processing

Brain dissections were performed under a stereoscope with regards to major sulci to identify cortical regions. Importantly, all dissections were performed by the same individual (T.J.N) to enable reproducibility and comparison between samples. Tissue was incubated in 4 ml of papain/DNAse solution (Worthington) for 20 min at 37C, after which it was carefully triturated with a glass pipette, filtered through a 40um cell strainer and washed with HBSS. The GW22 and GW25 samples were additionally passed through an ovomucoid gradient (Worthington) in order to minimize myelin debris in the captures. The final single-cell suspension was loaded onto a droplet-based library prep platform Chromium (10X Genomics) according to the manufacturer’s instructions. Version 2 was used for all samples except for GW19_2, GW16, and GW18_2 for which version 3 chemistry was utilized. cDNA libraries were quantified using an Agilent 2100 Bioanalyzer, and sequenced with an Illumina NovaSeq S4.

### Quality Control and Filtering

We filtered cells using highly stringent QC metrics. Briefly, we discarded potential doublets using the R package *scrublet* (Wolock, et al; PMID: 30954476) for each individual capture lane, then required at least 750 genes per cell and removed cells with high levels (>10%) of mitochondrial gene content. These strict metrics for quality control preserved no more than 40% of cells for downstream analysis, and re-analysis of the data for specific brain structures or cell types may benefit from less stringent QC for additional discovery. Our goal was to obtain clean populations with a high validation rate for a better understanding of arealization signatures. The resulting ∼700,000 cells passing all thresholds were used in downstream analyses.

### Clustering strategy

We used a recursive clustering workflow to understand the cell types present in our dataset. In order to minimize potential batch effects and to increase detection sensitivity of potential rare cell populations, we performed Louvain-Jaccard clustering on each individual sample first. After initial cell type classification, we sub-clustered all the cells belonging to a cell type to generate the most granular cell subtypes possible. We then correlated subtypes between individuals based upon the gene scores in all marker genes to bridge any batch effects, and iteratively combined clusters across all individuals and cell types. For this study, we combined the clusters within a single cell type across all individuals once, and again with all clusters from all individuals and cell types, resulting in two iterative combinations. The annotations at each step are preserved in the supplementary tables to enable reconstruction at any point in the pipeline.

### Hierarchical Clustering of Clusters

Cluster hierarchies are generated from matrices correlating all clusters to one another using Pearson’s correlation in the space of gene scores for all marker genes across all groups. Hierarchical clustering is performed within Morpheus (https://software.broadinstitute.org/morpheus) across all rows and columns using one minus the Pearson correlation for the distance metric.

### Constellation plots

To visualize and quantify the global relationships and connectedness between cell types, cell type subclusters, or cell type-area groups, we implemented the constellation plots described in Tasic, 2018^2^, by adapting the code made available at https://github.com/AllenInstitute/scrattch.hicat/. Briefly, we represented each group of cells as a node, whose size is proportional to the number of cells contained within it. Each node is positioned at the centroid of the UMAP coordinates of its cells. Edges represent relationships between nodes, and were calculated by obtaining the 15 nearest neighbors for each cell in PCA space (PCs 1:50), then determining, at each cluster, the fraction of neighbors belonging to a different cluster. An edge is drawn between 2 nodes if >5% of nearest neighbors belong to the opposite cluster in at least one of them. An edge’s width at the node is proportional to the fraction of nearest neighbors belonging to the opposite node, with the maximum fraction of out-of-node neighbors across all clusters represented as an edge width of 100% and equal to node width. The full code adaptation and implementation of this analysis is described in the function *buildConstellationPlot* in this paper’s Github repository.

### Quantification of Constellation Plots

Constellation plots were quantified by using a summary of the input values described above. For each cell type or area connection, the number of edges between two groups was multiplied by the average fraction of cells meeting the threshold for a connection within the group. This resulting number was called the connectivity index.

### Module Eigengene Calculations

Module eigengenes were calculated for numerous gene sets using the the R package WGCNA^35^. Scores were generated for each set of up to 10,000 randomly subsetted cells from the group using the function moduleEigengene function, Scores were calculated based on the intersection of the gene set of interest and genes expressed in the subset of cells. For the area-specific signatures, differential expression was performed as described above, and the gene signatures from late stage neurons across all areas were used to calculate module eigengenes for the radial glia and intermediate progenitor populations.

### Area-specific markers / Gene Score Calculations

The expression profiles of cells from each subcluster or cortical area were compared to those of all other cells using the two-sided Wilcoxon rank-sum test for differential gene expression implemented by the function FindAllMarkers in the R package Seurat and selected based on an adjusted p-value cutoff of 0.05. Adjusted p-values were based on Bonferroni correction using all features in the dataset. We performed this step separately for each cell type and each individual, since we observed that gene specificity was highly dynamic throughout the developmental process. We then combined the individual gene lists of each cell type and area, and annotated the stage(s) at which each gene appeared to be specific. We binned individuals into three stages: early (GW14, 16 and 17), middle (GW18, 19, 20), and late (GW22, GW25). We ranked upregulated genes by specificity by calculating their gene score, which we defined as the result of a gene’s average log fold-change x its enrichment ratio, in turn defined as the percentage of cells expressing the gene in the cluster of interest / percentage of cells expressing in the complement of all cells. Dot Plots used to visualize the expression of distinct marker genes across cell types and/or cortical areas were generated the custom function makeDotPlot available in our code repository, which makes use of the Seurat function DotPlot. Briefly, for each gene, the average expression value of all non-zero cells from each group (cortical area) is scaled using the base R function scale(), yielding obtaining z-scores. Scaling is done in order to enable the visualization of genes across vastly different expression ranges on the same color scale.

### Transcription Factor Annotation

Areally enriched marker genes obtained as described above were annotated against a comprehensive list of 1,632 human transcription factors described in^36^ and downloaded from the transcription factor database AnimalTFDB3^37^, available at http://bioinfo.life.hust.edu.cn/static/AnimalTFDB3/download/Homo_sapiens_TF.

### Gene signature overlap and Sankey Diagrams

In order to quantify the degree of (dis)similarity of molecular signatures across distinct cell types, cortical areas, and/or developmental stages, we calculated the overlap between all sets of cell type and area-specific gene markers at each stage, and visualized these comparisons using Sankey diagrams using the function *ggSankey* from the *ggv*is R package. We then calculated the proportion of genes for each node shared with every other node, and clustered nodes hierarchically by calculating their euclidean distances based on their proportions of shared genes. The code used to construct the overlap matrices, create the plots and quantify the results is described in the functions *buildSankey* and *buildHeatmap* in our Github repository.

### RNA Velocity

Velocity estimates were calculated using the Python 3 packages Velocyto v0.17^20^ and scVelo v0.2.2^21^. Reads that passed quality control after clustering were used as input for the *Velocyto* command line implementation. The human expressed repeat annotation file was retrieved from the UCSC genome browser. The genome annotation file used was provided by *CellRanger*. The output loom files were merged and used in scVelo to estimate velocity. For the combined cortical analysis, cells underwent randomized subsampling (fraction=0.5), and were filtered based on the following parameters: minimum total counts = 200, minimum spliced counts = 20 and minimum unspliced counts = 10. The final processed object generated a new UMI count matrix of 18,970 genes across 195,775 cells, for which the velocity embedding was estimated using the stochastic model. The embedding was visualized using Uniform Manifold and Approximation and Projection of dimension reduction. The velocity genes were matched by cortical area and were estimated using the rank velocity genes function in scVelo. Computational analysis of the transcriptomic data described in detail above were performed using *R 4.0*^*38*^ and Python 3, the R packages *Seurat* (version 2 and version 3)^39, 40^, *googleVis*^*41*^, *dplyr and ggplot2*^*42*^, the Python packages *Velocyto v0.17*^*20*^ and *scVelo v0.2.2*^*21*^ as well as the custom-built R functions described. Our reproducible code is available in the Github repository associated with this manuscript.

### Validation Marker Gene Selection

Marker genes for validation with the spatial omics platform were chosen first by identify useful cell type markers within the dataset. *SOX2* was chosen to mark radial glia, *EOMES* was chosen to mark IPCs, and *BCL11B* and *SATB2* were chosen to marker excitatory neuronal populations with previously validated changing co-expression patterns. *POLR2A* was used as a positive control for the technology. The remaining genes were selected based upon their status as a specific marker gene for excitatory neuron clusters of interest.

### Rebus Esper Spatial Omics Platform

Samples for spatial transcriptomics were dissected from primary tissue as described above. Samples were flash frozen in OCT following the protocol described in the osmFISH protocol^43^. Samples were then mounted to APS coated coverslips, and fixed for 10 minutes in 4% PFA. Samples were then washed with PBS, and processed for spatial analysis. The spatially resolved, multiplexed in situ RNA detection and analysis was performed using the automated Rebus Esper spatial omics platform (Rebus Biosystems, Inc., Santa Clara, CA). The system integrates synthetic aperture optics (SAO) microscopy^44^, fluidics and image processing software and was used in conjunction with single-molecule fluorescence *in situ* hybridization (smFISH) chemistry. Individual transcripts from target genes were automatically detected, counted, and assigned to individual cells, generating a Cell x Feature matrix that contains gene expression and spatial location data for each individual cell, as well as registered imaging data, as follows:

Rebus Biosystems proprietary software was used to design primary target probes (22-96 oligos) and corresponding unique readout probes (assigned and labeled with Atto dyes) for each gene. The oligos were purchased from Integrated DNA Technologies and resuspended at 100µM in TE buffer. Coverslips (24 × 60 mm, No. 1.5, Cat. # 1152460, Azer Scientific) were functionalized as previously published^43^. Fresh frozen brain tissue sections (10 µm) were cut on a cryostat, mounted on the treated coverslips and fixed for 10 min with 4% paraformaldehyde (Alfa Aesar, Cat#43368) in PBS at room temperature, rinsed twice with PBS at room temperature and stored in 70% ethanol at 4°Cbefore use. The sample section on the coverslip was assembled into a flow cell, which was then loaded onto the instrument. The hybridization cycles and imaging were done automatically under the instrumental control software. Briefly, primary probes for all target genes were initially hybridized for 6 hours and probes not specifically bound were washed away. Readout probes labeled with Atto532, Atto594 and Atto647N dyes for the first 3 genes were then hybridized, washed, counterstained with DAPI and then imaged with an Andor sCMOS camera (Zyla 4.2 Plus, Oxford Instruments) through 20xC, 0.45NA dry lens (CFI S Plan Fluor ELWD, Nikon) with 365nm LED for DAPI and 532nm, 595nm and 647nm lasers configured for SAO imaging. Multiple fields of view (FOVs) were imaged for each channel within the region of interest (ROI). Single Z-planes with 2.8µm depth of field were acquired for each field of view. After imaging, the first 3 readout probes were stripped and the readout probes for the next 3 genes were then hybridized, imaged, and stripped. This process was repeated until readout was completed for all genes.

Using the Rebus Esper image processing software, the raw images were reconstructed to generate high-resolution images (equivalent or better than images obtained with a 100x oil immersion lens). RNA spots were automatically detected to generate high fidelity RNA spot tables containing xy positions and signal intensities. Nuclei segmentation software based on StarDist^45^ identified individual cells by finding nuclear boundaries from DAPI images. The detected RNA spots were then assigned to each cell using maximum distance thresholds. The resulting CellxFeature matrix contains gene counts per cell along with annotations for cell location and nuclear size.

### Kernel Density Estimation Plots

Kernel density estimation plots were created from individual gene spot location maps retrieved from the spatial transcriptomics pipeline. They were created using the seaborn kdeplot function in Python with shading and cmap coloring. They were merged together for Figure 4 with Adobe Illustrator’s overlay and darken features, using 50% opacity.

### Spatial Co-expression Analysis

Using the cell by feature matrices, we eliminated all spots with less than 10 counts for signal. Pearson’s correlations were then performed across the genes within each dataset and filtered for self-correlation. Positive control (POL2RA) and non-excitatory neuron cell type markers (SOX2, EOMES, DLX6) were removed from the analysis. Interactions of 0.05 or more were preserved and visualized with Cytoscape v3.8.2 using a force-directed biolayout. Individual nodes were colored by their color in the merged image file in Figure 4B.

## Supplementary Table Legends

**Supplementary Table 1**. Table of metadata for all cells from the whole brain analysis.

Supplementary Table 2. Cluster markers for the final iteration of whole brain combined clustering analysis. Markers were calculated using the Wilcoxon rank sum test. Table shows gene name, p-value, average log2 fold change, the percentage of cells within the cluster expressing the gene (pct.1), the percentage of cells outside the cluster expressing the gene (pct.2), and adjusted p-value. Positive log-fold change values indicate that the feature is more highly expressed in the cluster of interest.

**Supplementary Table 3**. Region specific marker genes, including those at the cell type specific level. Markers were calculated using the Wilcoxon rank sum test. Table shows gene name, p-value, average log_2_ fold change, the percentage of cells within the cluster expressing the gene (pct.1), the percentage of cells outside the cluster expressing the gene (pct.2), and adjusted p-value. Positive log-fold change values indicate that the feature is more highly expressed in the cluster of interest.

**Supplementary Table 4**. Cell type specific differential expression across the neocortex, allocortex (hippocampus), and proneocortex (cingulate). Markers were calculated using the Wilcoxon rank sum test. Table shows gene name, p-value, average log_2_ fold change, the percentage of cells within the cluster expressing the gene (pct.1), the percentage of cells outside the cluster expressing the gene (pct.2), and adjusted p-value. Positive log-fold change values indicate that the feature is more highly expressed in the cluster of interest.

**Supplementary Table 5**. Table of metadata for all neocortex cells.

**Supplementary Table 6**. Cluster markers for the final iteration of the neocortex combined clustering analysis. Markers were calculated using the Wilcoxon rank sum test. Table shows gene name, p-value, average log_2_ fold change, the percentage of cells within the cluster expressing the gene (pct.1), the percentage of cells outside the cluster expressing the gene (pct.2), and adjusted p-value.

**Supplementary Table 7**. Broader cell type signatures for radial glia, IPCs, and excitatory neurons in the Neu

**Supplementary Table 8**. All differentially expressed genes by cell type across all cortical areas. Markers were calculated using the Wilcoxon rank sum test. Table shows gene name, p-value, average log_2_ fold change, the percentage of cells within the cluster expressing the gene (pct.1), the percentage of cells outside the cluster expressing the gene (pct.2), and adjusted p-value.

**Supplementary Table 9**. CellxFeature matrix for all spots assigned to cells for the PFC slice analysis, includes x-y coordinates of the GW20 tissue.

**Supplementary Table 10**. CellxFeature matrix for all spots assigned to cells for the somatosensory cortex slice analysis, includes x-y coordinates of the GW20 tissue.

**Supplementary Table 11**. CellxFeature matrix for all spots assigned to cells for the temporal cortex slice analysis, includes x-y coordinates of the GW20 tissue.

**Supplementary Table 12**. CellxFeature matrix for all spots assigned to cells for the V1 cortex slice analysis, includes x-y coordinates of the GW20 tissue.

**Supplementary Table 13**. CellxFeature matrix for all spots assigned to cells for the PFC slice analysis, includes x-y coordinates of the GW16 tissue.

**Supplementary Table 14**. CellxFeature matrix for all spots assigned to cells for the somatosensory cortex slice analysis, includes x-y coordinates of the GW16 tissue.

**Supplementary Table 15**. CellxFeature matrix for all spots assigned to cells for the temporal cortex slice analysis, includes x-y coordinates of the GW16 tissue.

**Supplementary Table 16**. CellxFeature matrix for all spots assigned to cells for the V1 cortex slice analysis, includes x-y coordinates of the GW16 tissue.

