## supplementary_figures for "An Atlas of Cortical Arealization Identifies Dynamic Molecular Signatures"

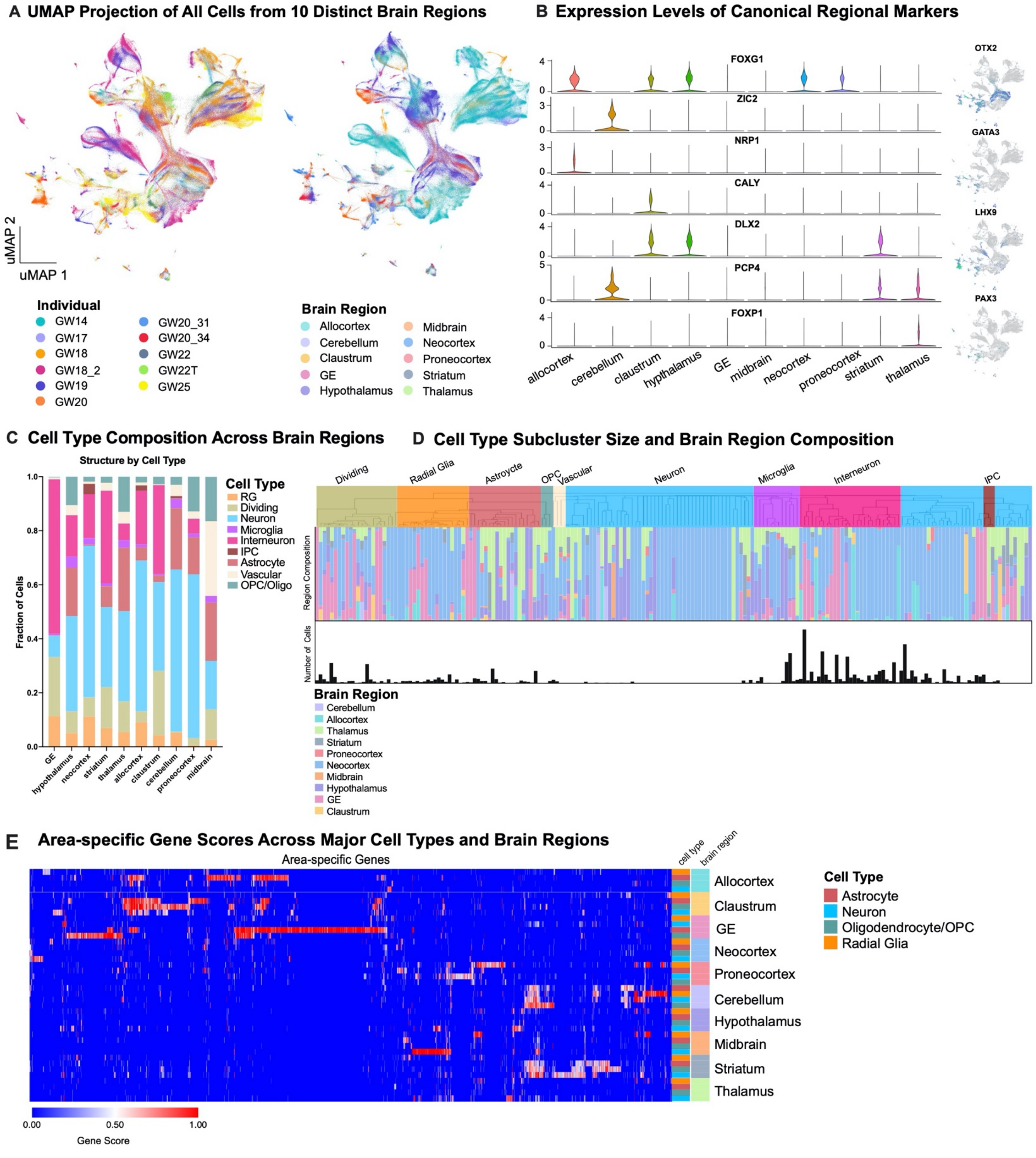


**Supplementary Figure 1. Single-cell analysis across the whole developing human brain**
**a)** UMAP plots showing the representation of samples by individual and brain region. There is strong intermixing across individuals, but more segregation by stage. **b)** On the left, violin plots of known and novel brain region-enriched marker genes. On the right are feature plots of cell type specific transcription factors. **c)** Histogram depicting the cell type composition as determined by the single-cell analysis for each sampled brain region, showing similar distributions across regions, but with known enrichments for inhibitory interneurons in the ganglionic eminences (GE), and other small enrichments for specific cell types in other regions. **d)** Hierarchical clustering of 210 neocortex clusters based upon Pearson correlations across cluster markers.

Each bar is colored based upon the major cell type assigned to that cluster. Beneath the clusters are histograms showing the fraction of cells from each area contributing to the cluster, and below that is a barchart showing the relative number of cells in each cluster (log2 transformed numbers ranging from 0 to 20). **e)** Heatmap representing the universe of area-specific genes for each cell type. Gene score, a metric that combines specificity and fold change, is shown from blue to red. Rows are grouped by brain region, and reveal that in many structures, area-specific genes cross cell types.

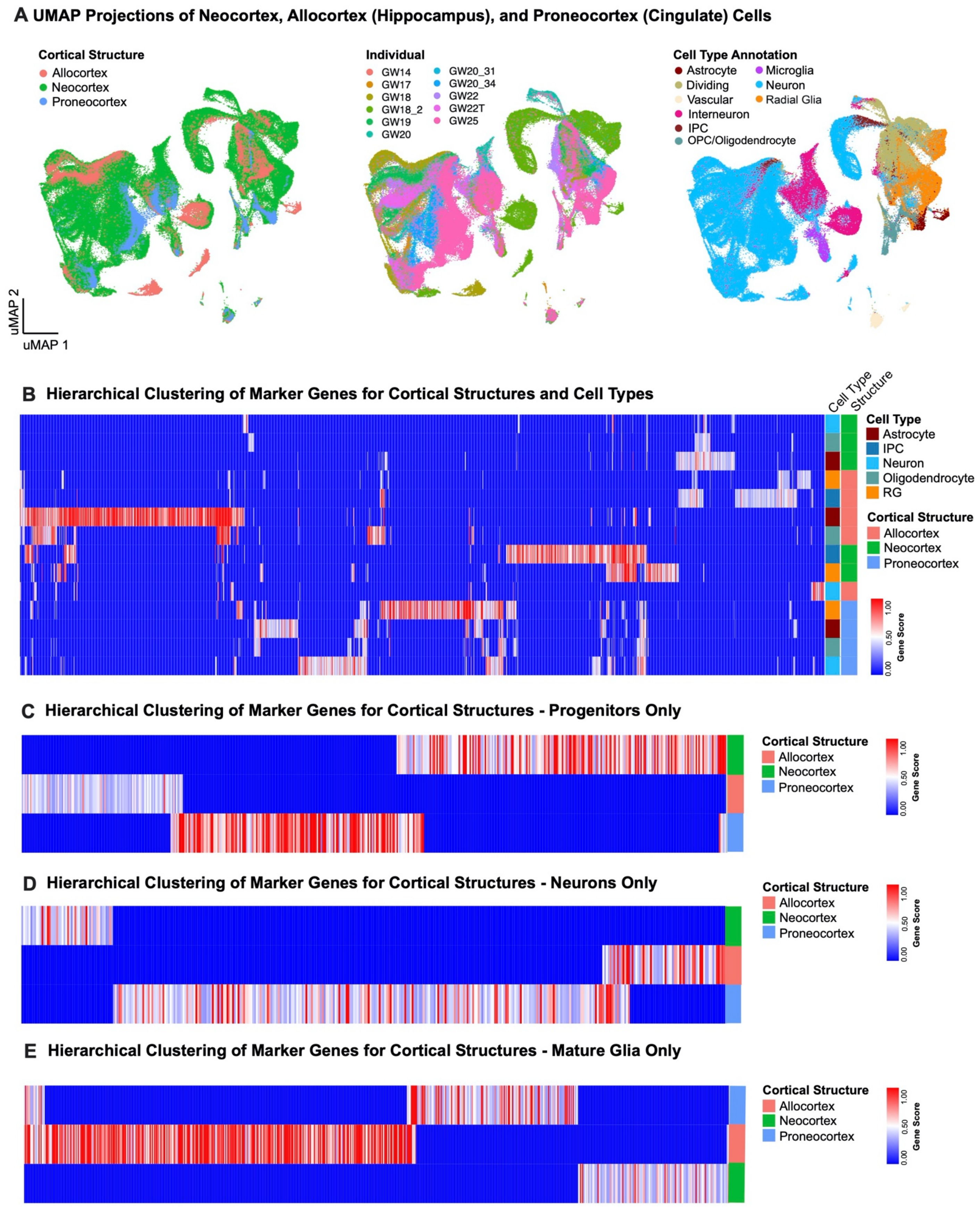


**Supplementary Figure 2. Single-cell analysis of regional identities across distinct cortical structures.**

**a)** UMAP projection of neocortex, allocortex and proneocortex cells alone, using cell type annotations from the previous analysis. Left UMAP plot shows integration of neocortex and allocortex cells (green and salmon, respectively), but exclusion of proneocortex cells (blue). In the middle UMAP plot, cells are colored by individual. Right UMAP plot shows the distribution and strong segregation by cell type. **b)** Marker genes (columns) specific to distinct cortical regions by cell type. Rows and columns are hierarchically clustered columns using one minus Pearson correlation for the distance metric. Heatmap reflects strong regional transcriptional identities, even among these 3 cortical structures. Additionally, it highlights expression signatures that cross cell type boundaries. **c)** Marker genes (columns) specific to distinct cortical regions analyzed in progenitors only. Rows and columns are hierarchically clustered columns using one minus Pearson correlation for the distance metric. **d)** Marker genes (columns) specific to distinct cortical regions analyzed in excitatory neurons only. Rows and columns are hierarchically clustered columns using one minus Pearson correlation for the distance metric. **e)** Marker genes (columns) specific to distinct cortical regions analyzed in mature glia only. Rows and columns are hierarchically clustered columns using one minus Pearson correlation for the distance metric.

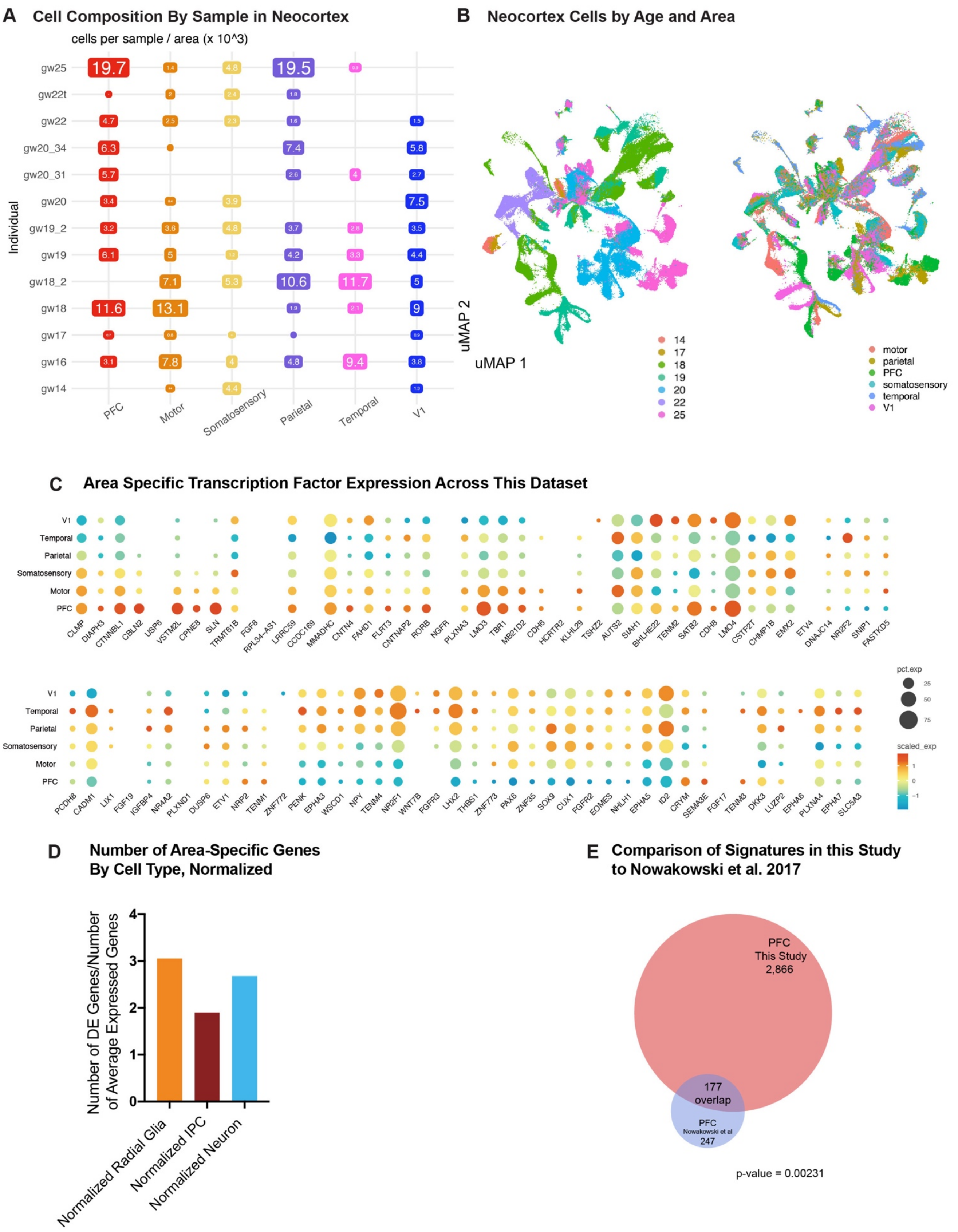


**Supplementary Figure 3. Single-cell analysis across the developing human neocortex**

**a)** Matrix showing the distribution of the number of cells across areal dissections for each of the individuals samples. Number of cells is shown in boxes sized proportionally to the number of cells from that individual and region, times 1000. **b)** UMAP plots showing the representation of samples by age (GW) (left) and brain region (right). There is strong intermixing across individuals, but more segregation by stage. **c)** Dot plot of area-specific genes as identified in Cadwell, et al 2019 as they are expressed in our dataset. Most genes show expected expression patterns, though some deviate from these expectations, likely because many of these genes have been characterized in the rodent. **d)** Quantification of the number of differentially expressed areal genes from each cell type, using a union of all genes as calculated across each individual in the dataset normalized by the average number of genes expressed within that cell type. **e)** Venn diagram showing the overlap between the PFC/V1 genes in the Nowakowski, et al 2017 dataset and this study. p-value = 0.00231, Chi-square test.

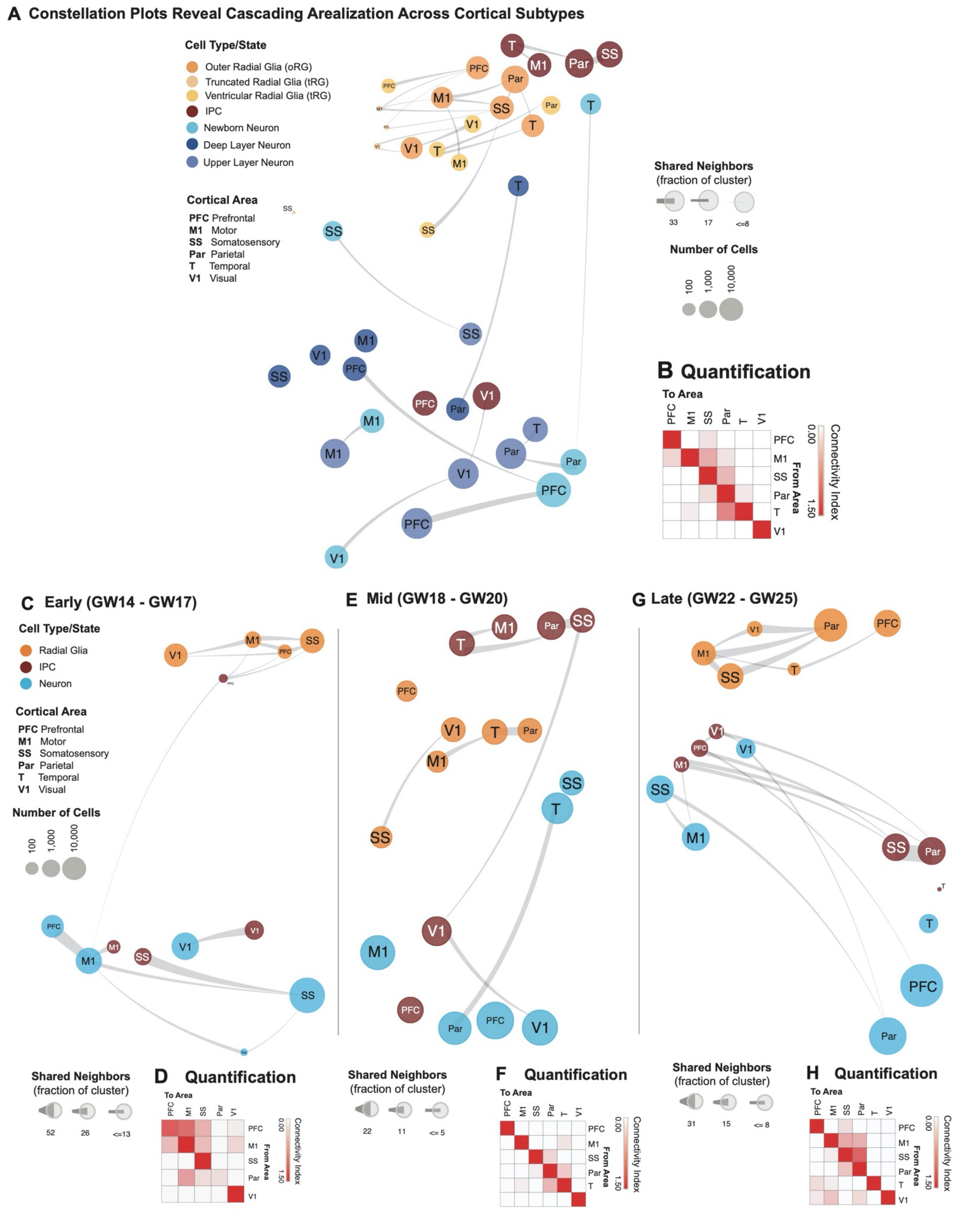


**Supplementary Figure 4. Constellation plots of cortical areas across cell subtypes and developmental stages a)** Constellation plots of excitatory lineage grouped by cortical area and annotated by cell subtype highlight cascading differences in areal identity, with similarities between cell types from the same region. Each dot is scaled proportionally to the number of cells represented by that analysis. The thickness of the connecting line on each end represents the fraction of cells within each group with neighbors in connected groups. Dot color represents cell type while text over the dot marks cortical area. **b)** Quantification of the constellation plots, with 'towards area' in columns and 'from area' in rows. The connectivity index from white to red integrates the number of connections between two cell types as well as the average fraction of cells from each cluster contributing to each connection. **c)** Constellation plots of excitatory lineage grouped by cortical area within early (GW14 – GW17) samples. **d)** Quantification of the early constellation plots. **e)** Constellation plots of excitatory lineage grouped by cortical area within mid-stage (GW18 – GW20) samples. **f)** Quantification of the mid-stage constellation plots. **g)** Constellation plots of excitatory lineage grouped by cortical area within late-stage (GW18 – GW20) samples. **h)** Quantification of the late-stage constellation plots.

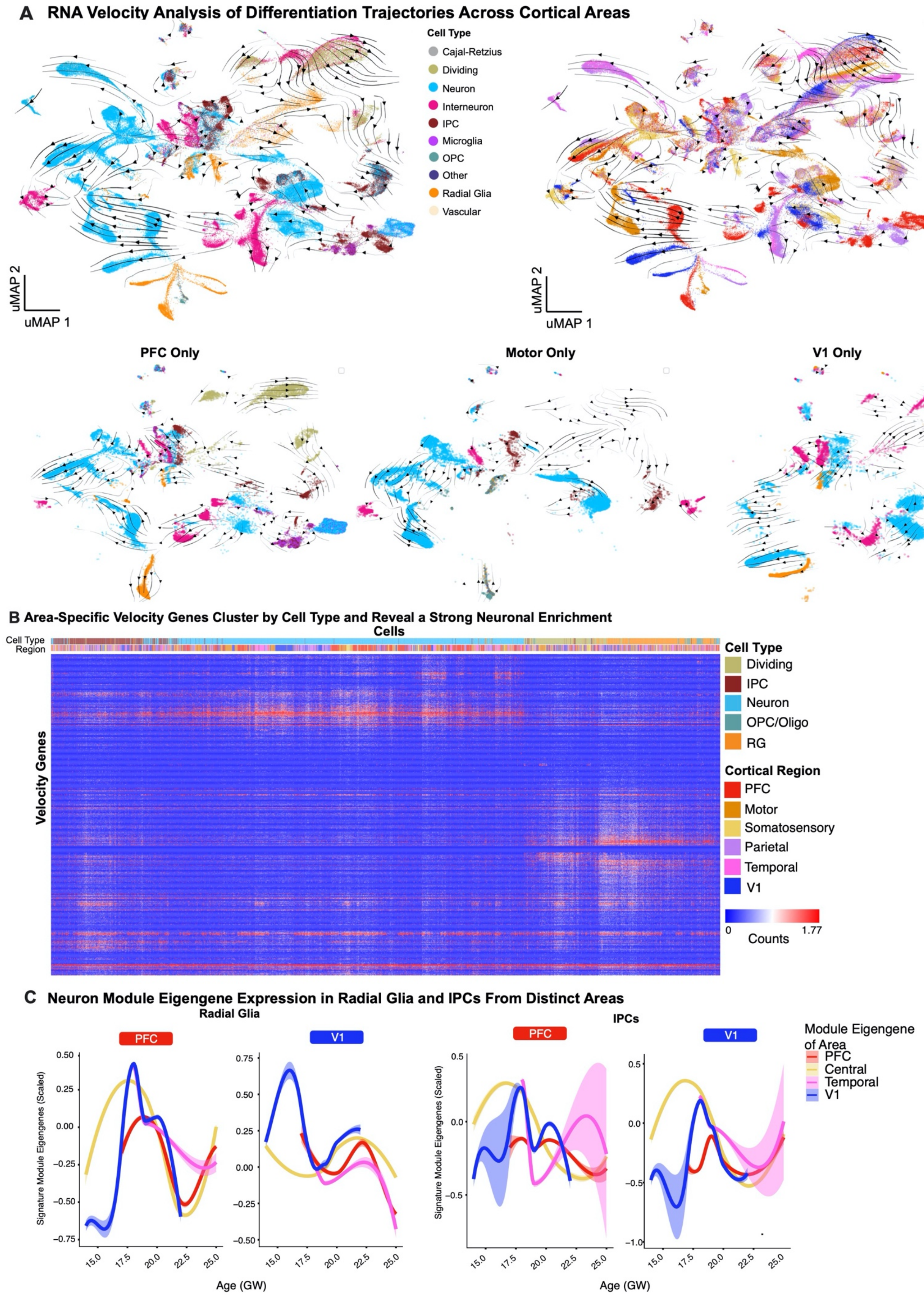


**Supplementary Figure 5. Trajectories of differentiation and changes in gene signatures.**

**a)** RNA velocity analysis was performed using all necortical cells from the dataset. Stream arrows depict predicted differentiation trajectories. Cells are shown in UMAP space, and are colored by cell type (left panel) and by cortical area (right panel). In the second row, zoomed in streams are shown for individual cortical areas PFC, motor and V1 only colored by cell type. **b)** Normalized counts for area-specific genes (rows) that were detected by the RNA-velocity algorithm are shown across a random subset of 20,000 cells (columns). Rows and columns were hierarchically clustered using a one minus pearson correlation distance matrix. Cell type and region are shown for each cell (top color bar), and highlight cortical area specific excitatory neuron populations. **c)** Module eigengenes were calculated for radial glia (left) and IPCs (right) based upon the area specific gene signatures identified in neurons of that area. For both sets of plots, the gene expression signature of PFC neurons is shown on the left and the V1 signature is shown on the right. Lines indicate the signature strength for progenitors of each region across the timepoints sampled. Early in development, PFC radial glia are defined by a low V1 signature strength, while V1 radial glia are defined by a strong V1 signature and minimal signal for all other areal signatures. IPCs from the PFC and V1 are generally defined by the lack of a strong signature for any of the areas calculated.

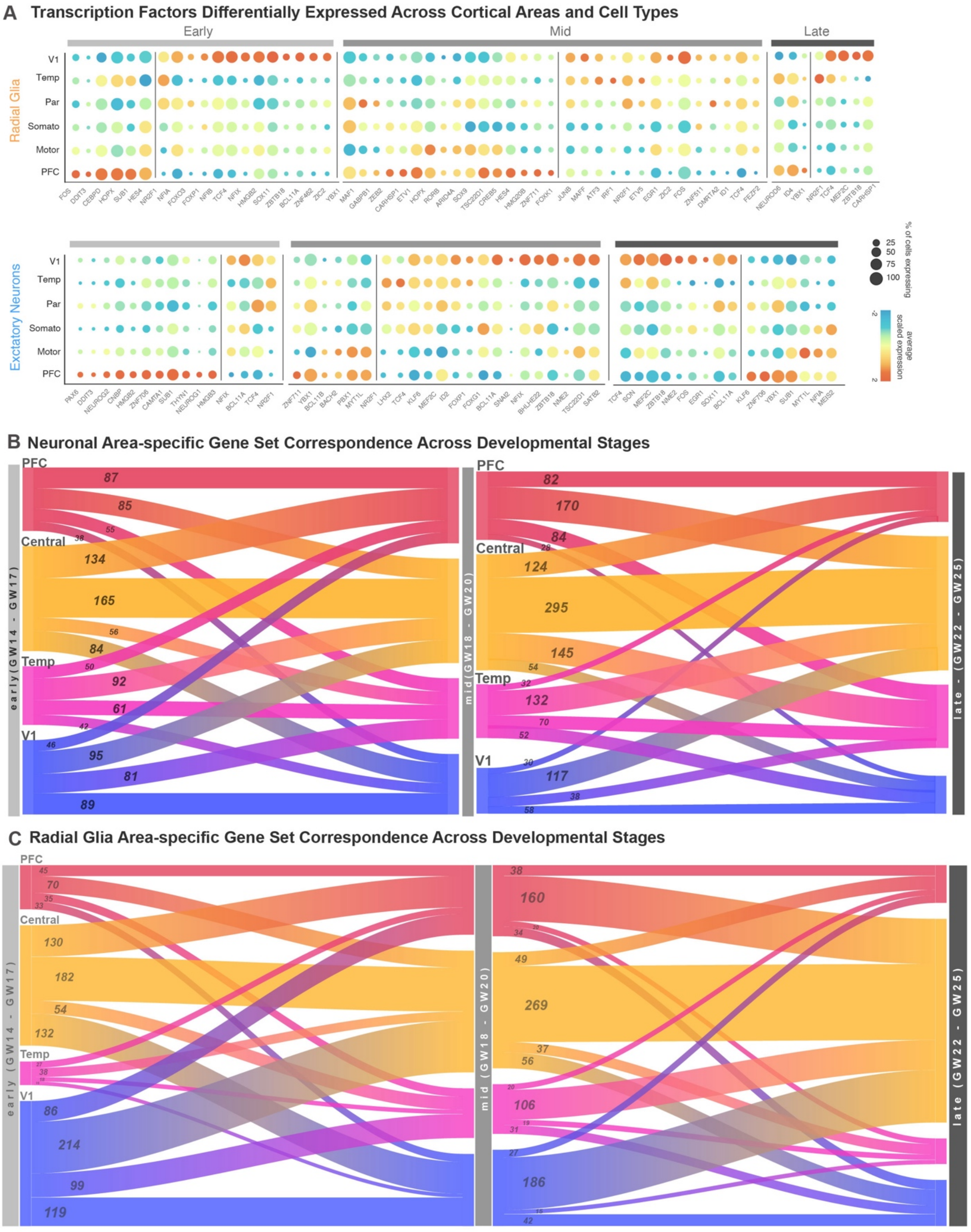


**Supplementary Figure 6. Gene Expression Dynamics Across Areas and Developmental Stages. a)** Dot plots transcription factors enriched across areas in radial glia (left) and excitatory neurons (right) relative to other cortical areas. Enrichment can occur via an increase in the number of cells expressing a given gene, an increase in the average expression level of expressing cells, or both. **b)** Sankey plots show the proportion of area-specific excitatory neuron genes shared across developmental stages. The number on each block line indicates how many genes are represented by that stream as the stream sizes are not to scale. The “central” area encompasses motor, parietal, and somatosensory areas. **c)** Sankey plots show the proportion of area-specific radial glia genes shared across developmental stages. The number on each block line indicates how many genes are represented by that stream as the stream sizes are not to scale.

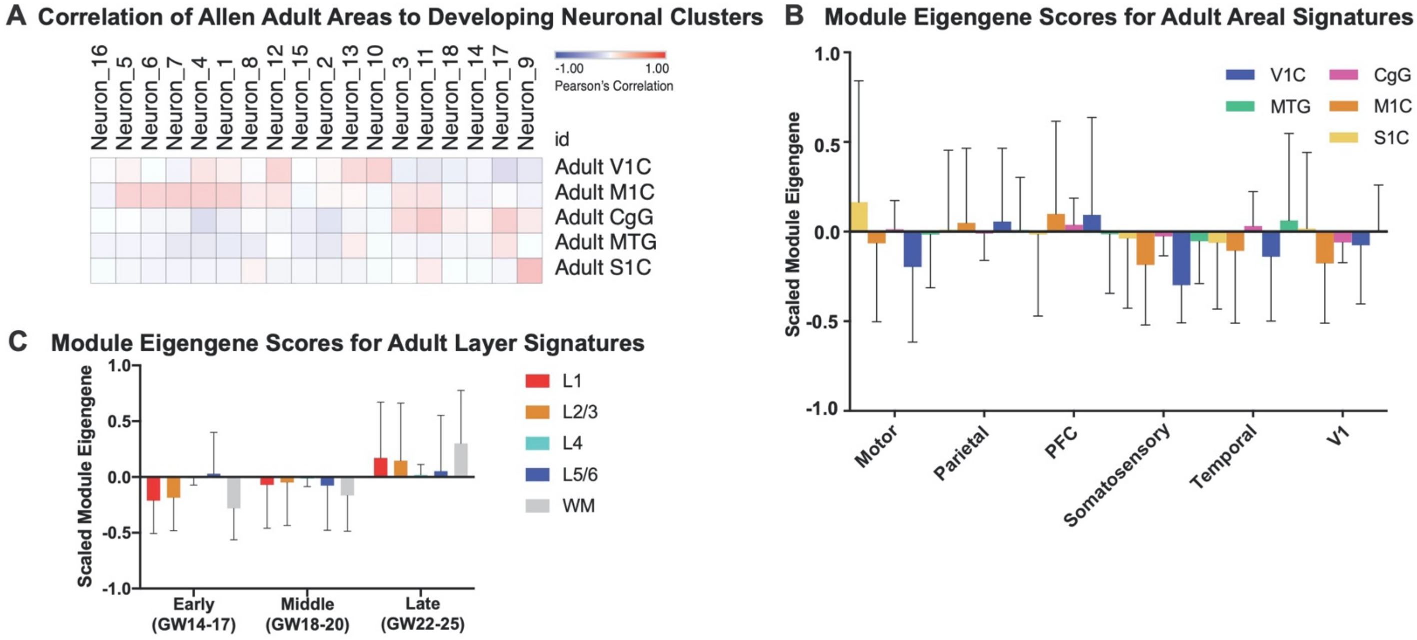


**Supplementary Figure 7. Signature Correspondence Between Developmental and Adult Samples**
**a)** Correlations were performed between the neuronal clusters in this dataset and the area specific identities calculated from adult human cortical areas as obtained from the Allen Institute Brain Map dataset. **b)** Module eigengene calculations of adult areal signatures show minimal to no correspondence to the signatures we identify in this study based upon low module eigengene values. **c)** Module eigengene values of Allen excitatory neurons layer specific signatures in each of the developmental stages within our dataset. Upper layer signatures emerge at late stages, while a small signal for deep layer identities can be observed at the earliest stages.

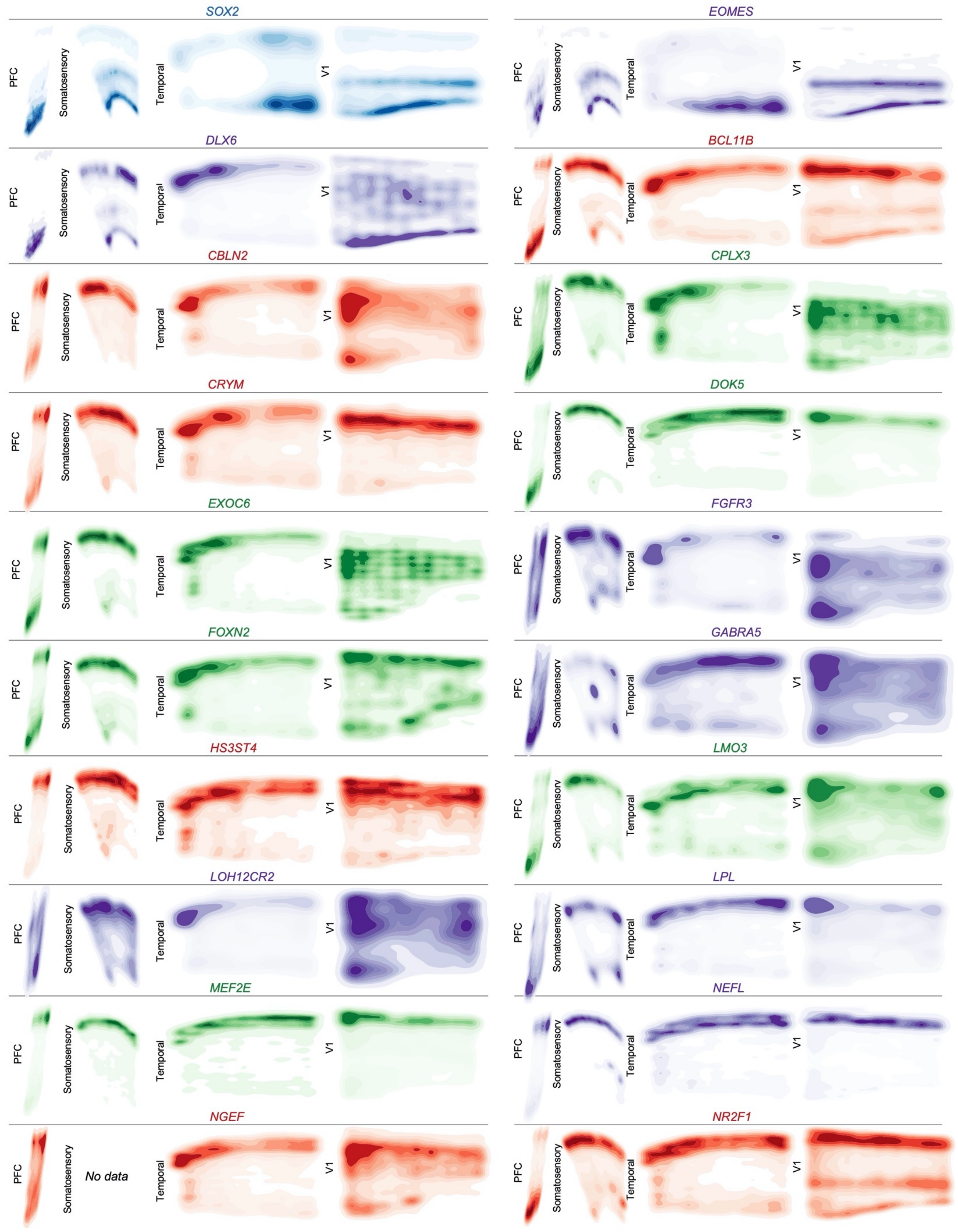


**Supplementary Figure 8. Spatial Expression Patterns of Cell Type and Neuronal Cluster Marker Genes**

**GW20 Part 1** Kernel density plots of each gene assayed using spatial RNA *in situ* analysis. Plots are made from all spots, and are shown across all four sampled cortical areas of a GW20 individual. Color is for emphasis of expression, but individual colors have no specific meaning.

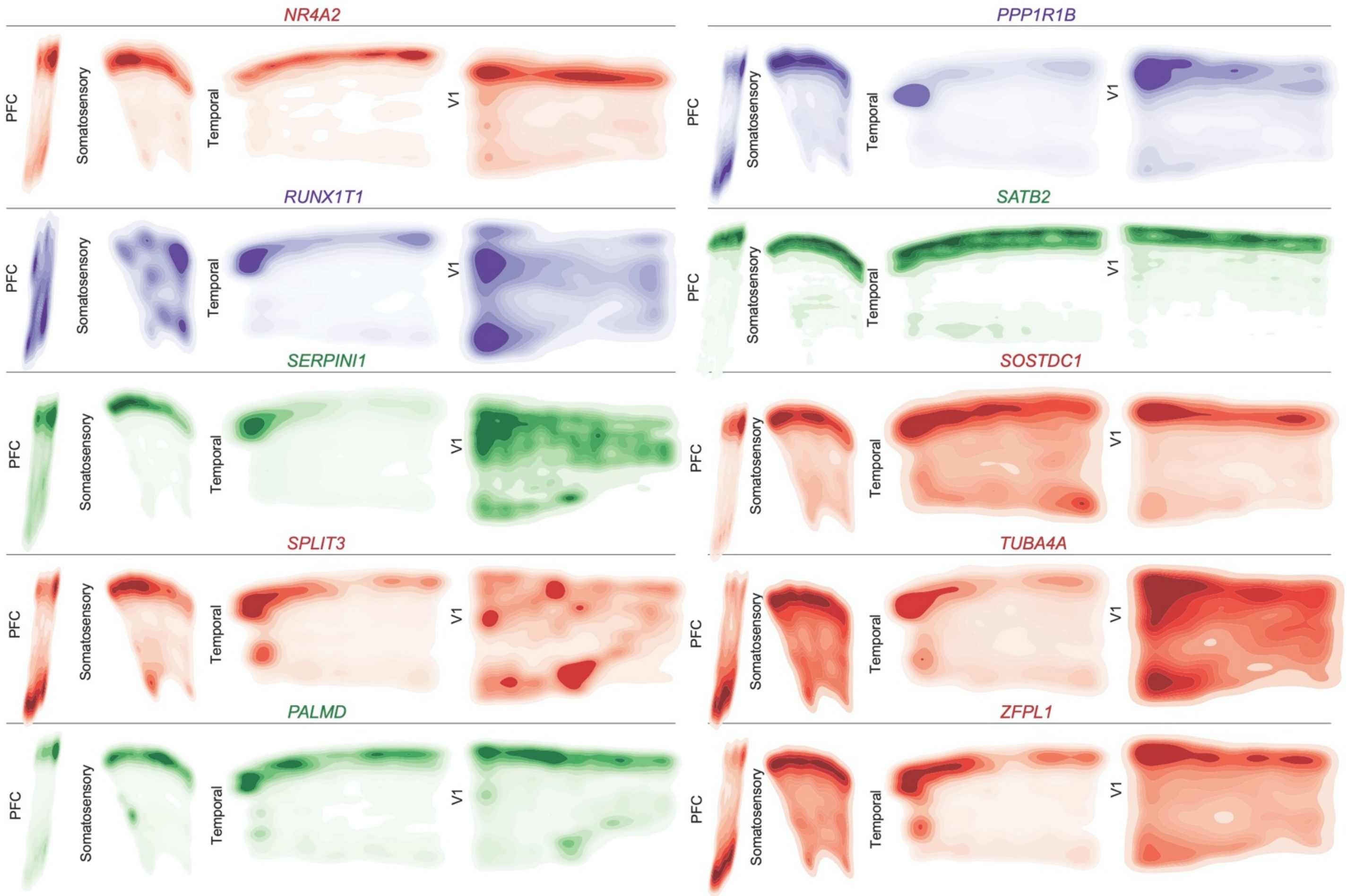


**Supplementary Figure 9. Spatial Expression Patterns of Cell Type and Neuronal Cluster Marker Genes GW20 Part 2** Kernel density plots of each gene assayed using spatial RNA *in situ* analysis. Plots are made from all spots, and are shown across all four sampled cortical areas of a GW20 individual. Color is for emphasis of expression, but individual colors have no specific meaning.

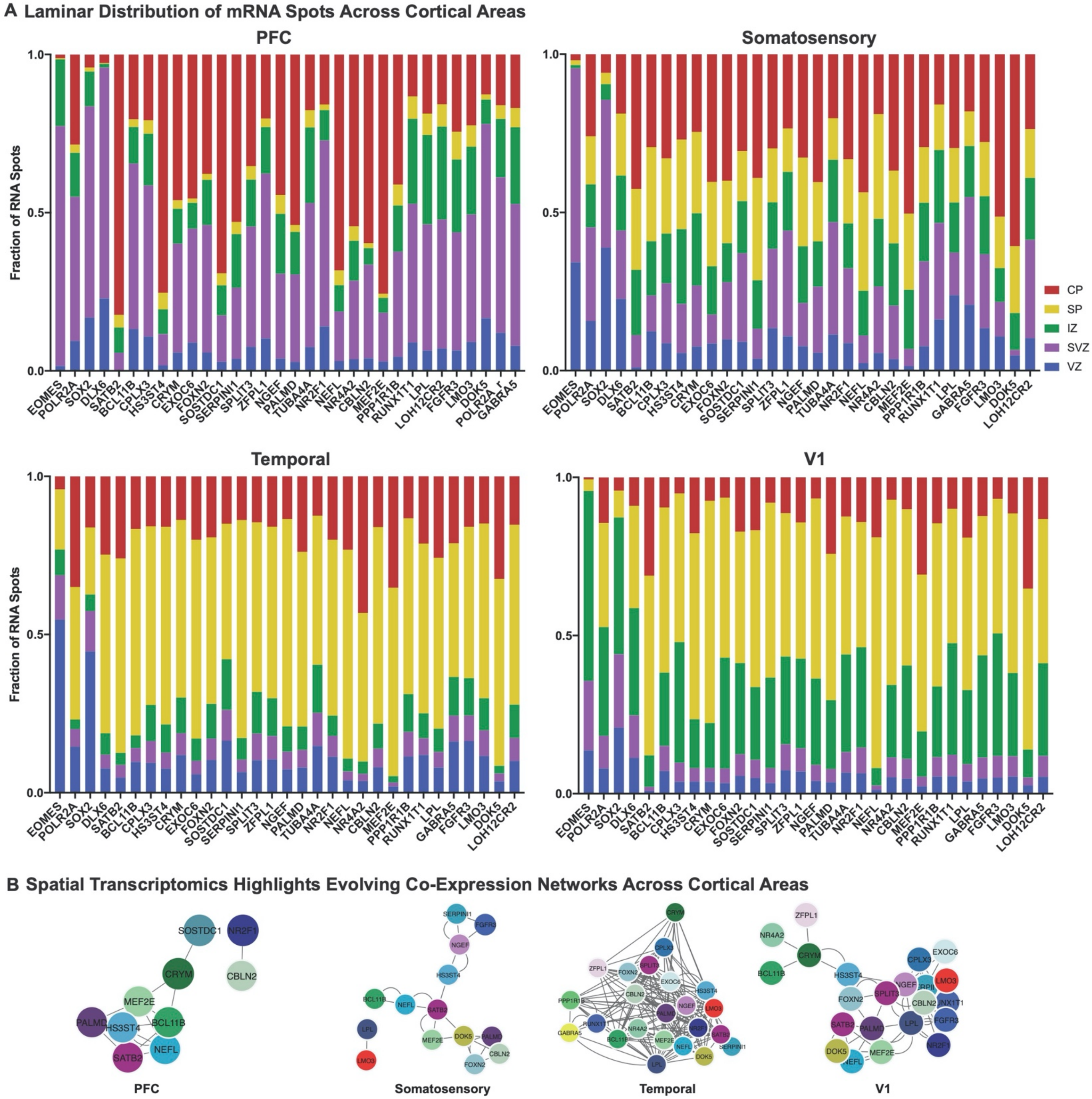


**Supplementary Figure 10. Laminar and network changes across cortical areas (GW20)**
**a)** We quantified the laminar distributions of each gene in each GW20 sample. The distributions are shown by laminar region, as annotated in Figure 4, and are represented as fraction of signal for each gene in each bin.
**b)** Using cell identities for individual gene spots, we computed co-expression networks for each sample to visualize changes in the co-expression patterns of the 31 genes analyzed across areas of the cortex. Gene pairs were classified as coexpressed if their Pearson correlation was >=0.05. Networks are shown in a force-directed layout reflective of interaction strength. Only neuronal marker genes are shown, colored as in panel (b).

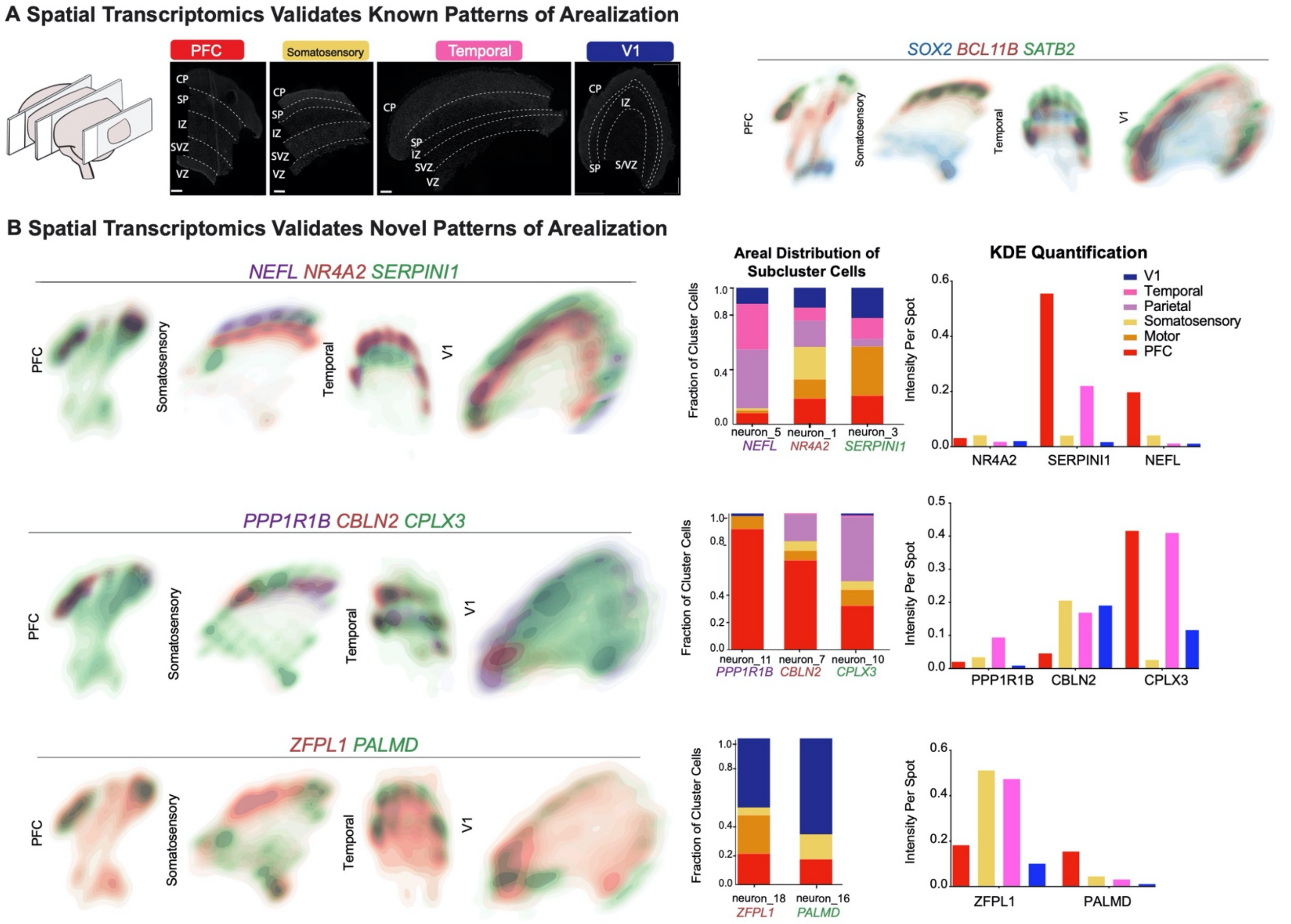


**Supplementary Figure 11. Spatial analysis of GW16 sample a)** Top left: Nucleus staining outlines tissue architecture, with the ventricular zone at the bottom and the cortical plate at the top. Top right: kernel density estimate (KDE) plots of positive control genes. SOX2 marks radial glia and the ventricular zone, while SATB2 and BCL11B mark the cortical plate. As previously described, SATB2 and BCL11B are co-expressed in frontal regions, but are mutually exclusive in occipital regions. Scale bar = 444 $\mu$M **b)** KDE plots of neuronal genes of interest. Genes were chosen as candidate markers for specific neuronal subclusters. Clusters being explored are named below the histogram each gene marker for this cluster is below its name. Stacked histograms show the expected ratio of clusters as a fraction of total composition. To the far right in each row, the quantification of the KDE plots is shown as intensity divided by the number of spots in order to reflect both the intensity of signal but also the pervasiveness of the marker to not artificially bias the analysis by examples of rare but intense signal. We see strong correspondence between the predicted spatial distribution of clusters and the signal in our spatial RNA analysis.

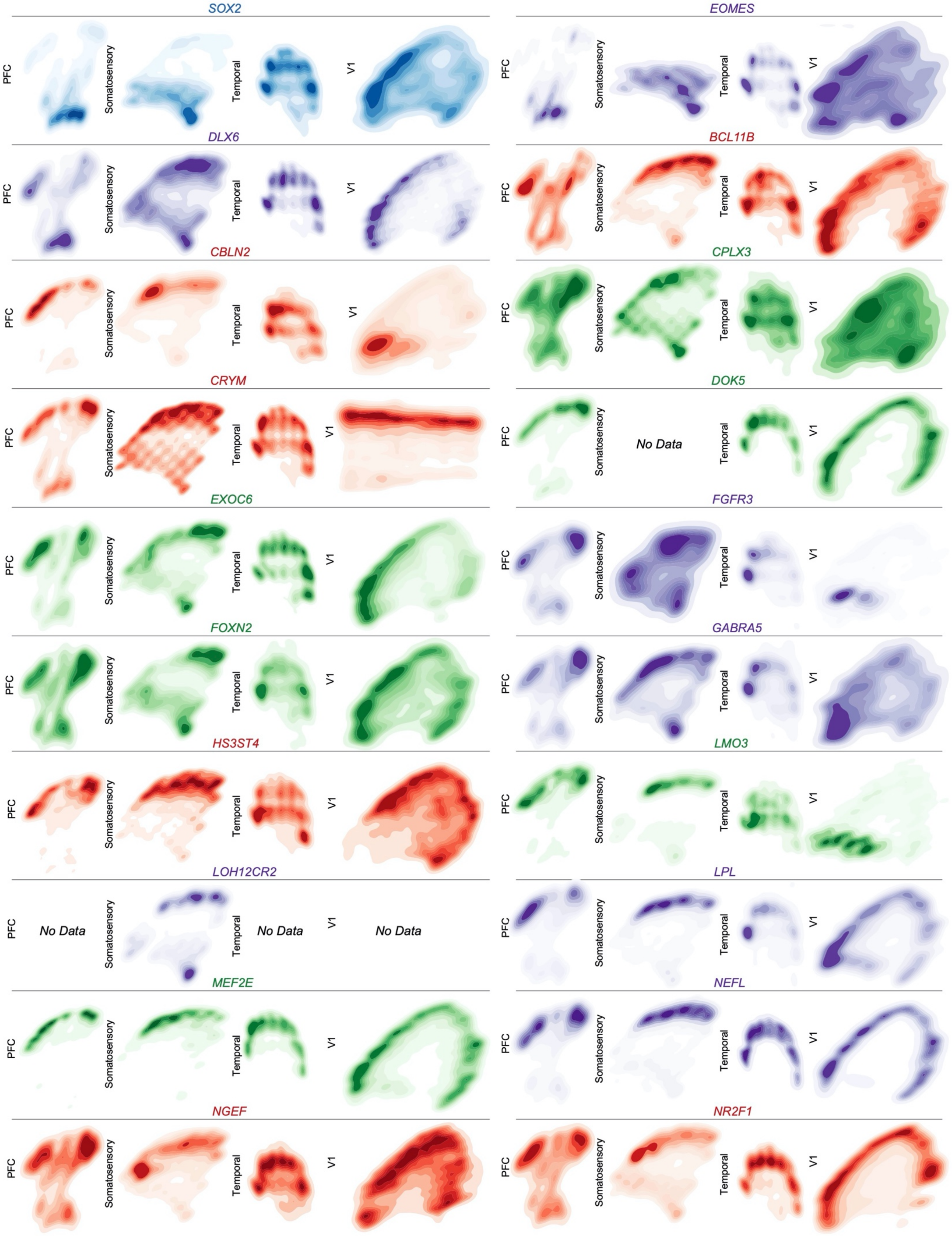


**Supplementary Figure 12. Spatial Expression Patterns of Cell Type and Neuronal Cluster Marker Genes**

**GW16 Part 1** Kernel density plots of each gene assayed using spatial RNA *in situ* analysis. Plots are made from all spots and are shown across all four sampled cortical areas of a GW16 individual. Color is for emphasis of expression, but individual colors have no specific meaning.

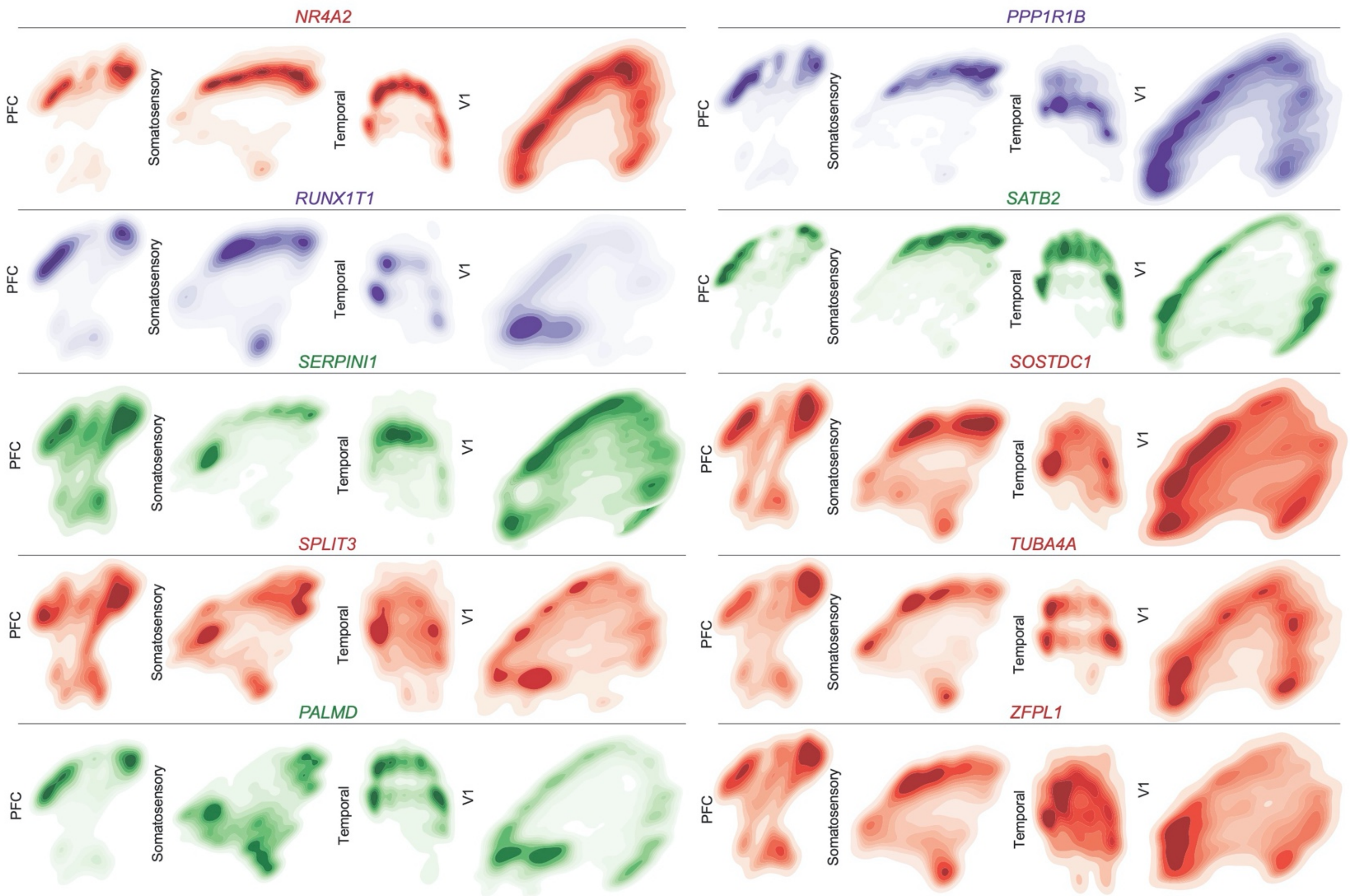


**Supplementary Figure 13. Spatial Expression Patterns of Cell Type and Neuronal Cluster Marker Genes GW16 Part 2** Kernel density plots of each gene assayed using spatial RNA *in situ* analysis. Plots are made from all spots, and are shown across all four sampled cortical areas of a GW16 individual. Color is for emphasis of expression, but individual colors have no specific meaning.

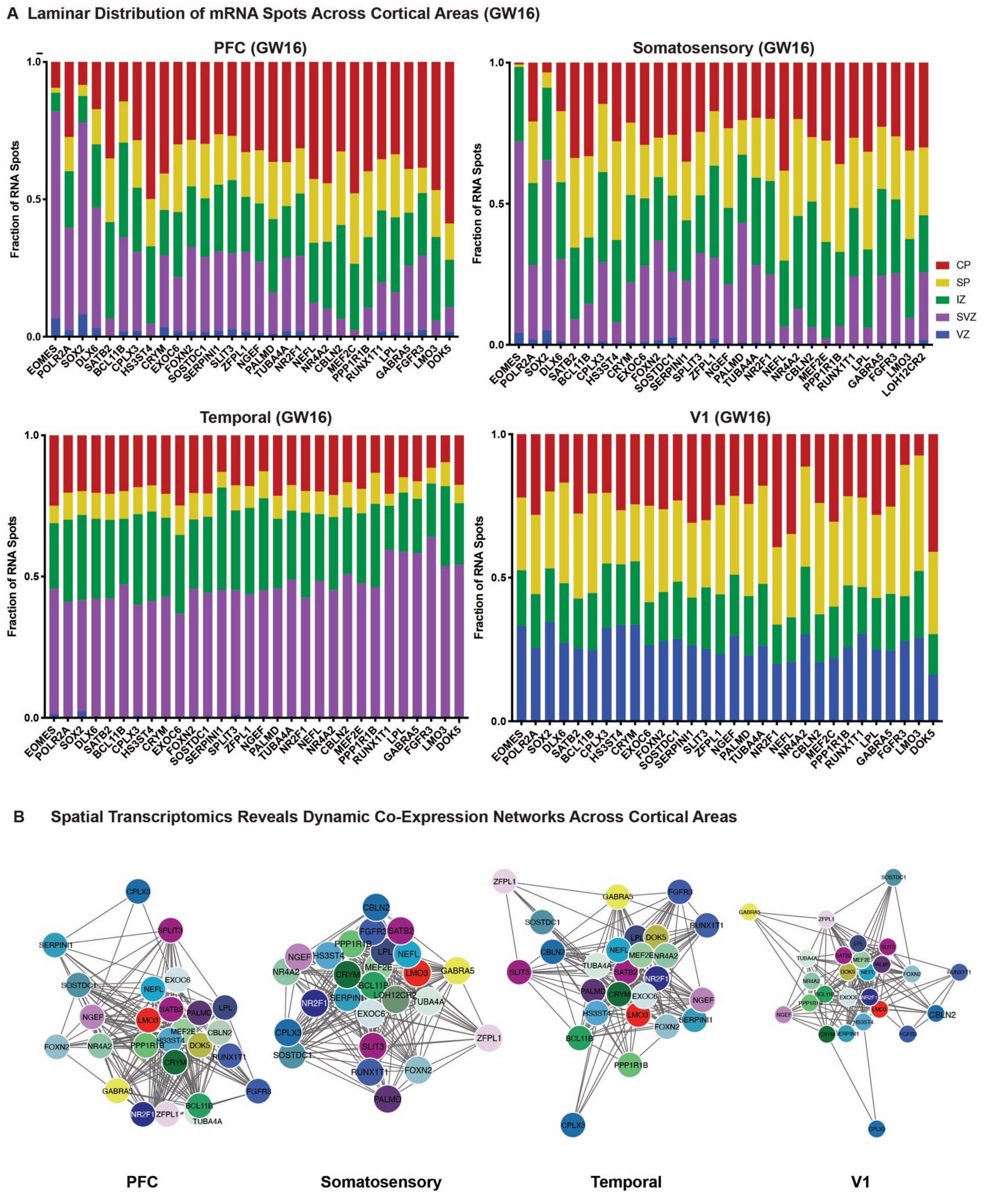
A Laminar Distribution of mRNA Spots Across Cortical Areas (GW16)
PFC (GW16)
Somatosensory (GW16)
Temporal (GW16)
V1 (GW16)
Fraction of RNA Spots
0.0
0.5
1.0
CP
SP
IZ
SVZ
VZ
EOMES
POLR2A
SOX2
DLX6
SATB2
BCL11B
CPLX3
HS3ST4
CRYM
EXOC6
FOXN2
SOSTDC1
SERPINI1
SLIT3
ZFPL1
NGEF
PALMD
TUBA4A
NR2F1
NEFL
NR4A2
CBLN2
MEF2C
PPP1R1B
RUNX1T1
LPL
GABRA5
FGFR3
LMO3
DOK5
LOH12CR2
B Spatial Transcriptomics Reveals Dynamic Co-Expression Networks Across Cortical Areas
PFC
Somatosensory
Temporal
V1

**Supplementary Figure 14. Laminar and network changes across cortical areas (GW16) a)** We quantified the laminar distributions of each gene in each GW16 sample. The distributions are shown by laminar region, as annotated in Figure 4, and are represented as fraction of signal for each gene in each bin. **b)** Using cell identities for individual gene spots, we computed co-expression networks for each sample to visualize changes in the co-expression patterns of the 31 genes analyzed across areas of the cortex. Gene pairs were classified as coexpressed if their Pearson correlation was >=0.05. Networks are shown in a force-directed layout reflective of interaction strength. Only neuronal marker genes are shown, colored as in panel (b).
